# Substrate Stiffness Dictates Unique Doxorubicin-induced Senescence-associated Secretory Phenotypes and Transcriptomic Signatures in Human Pulmonary Fibroblasts

**DOI:** 10.1101/2024.11.18.623471

**Authors:** Huixun Du, Jacob P. Rose, Joanna Bons, Li Guo, Taylor R. Valentino, Fei Wu, Jordan B. Burton, Nathan Basisty, Max Manwaring-Mueller, Priya Makhijani, Nan Chen, Veronica Chang, Shawn Winer, Judith Campisi, David Furman, Andras Nagy, Birgit Schilling, Daniel A. Winer

## Abstract

Cells are subjected to dynamic mechanical environments which impart forces and induce cellular responses. In age-related conditions like pulmonary fibrosis, there is both an increase in tissue stiffness and an accumulation of senescent cells. While senescent cells produce a senescence-associated secretory phenotype (SASP), the impact of physical stimuli on both cellular senescence and the SASP is not well understood. Here, we show that mechanical tension, modeled using cell culture substrate rigidity, influences senescent cell markers like SA-β-gal and secretory phenotypes. Comparing human primary pulmonary fibroblasts (IMR-90) cultured on physiological (2 kPa), fibrotic (50 kPa), and plastic (approximately 3 GPa) substrates, followed by senescence induction using doxorubicin, we identified unique high-stiffness-driven secretory protein profiles using mass spectrometry and transcriptomic signatures, both showing an enrichment in collagen proteins. Consistently, clusters of p21+ cells are seen in fibrotic regions of bleomycin induced pulmonary fibrosis in mice. Computational meta-analysis of single-cell RNA sequencing datasets from human interstitial lung disease confirmed these stiffness SASP genes are highly expressed in disease fibroblasts and strongly correlate with mechanotransduction and senescence-related pathways. Thus, mechanical forces shape cell senescence and their secretory phenotypes.

## Introduction

Cellular senescence is characterized by growth arrest, apoptosis resistance and altered gene expression that drives the development of a senescence-associated secretory phenotype (SASP). Senescent cells and their SASP are major contributors to aging and age-related diseases. Many age-related diseases are linked to fibroblasts and scar tissue formation, including pulmonary fibrotic diseases such as idiopathic pulmonary fibrosis, as well as fibrosis of other tissues like kidney, liver and fat, including obesity-related fibrosis [1,2]. Several stressors such as DNA damage, telomere instability, radiation and oncogene activation can induce cellular senescence [3]. Genotoxic compounds, like the anticancer drug doxorubicin, also induce senescence [4] and have been widely used to model senescence *in vitro* models [5].

As senescent cells accumulate with age, a chronic SASP has been considered a main driver of age-related pathologies [6]. The SASP includes a myriad of soluble cytokines, chemokines, growth factors and proteases released by senescent cells that trigger beneficial effects like wound healing, as well as deleterious effects such as inflammation and growth arrest in neighboring cells [7]. However, the SASP is heterogenous. Not only can the SASP be categorized into soluble SASP (sSASP), and extracellular vesicles SASP (eSASP), its identity varies widely depending on cell type and senescence trigger, making it a complex process to understand [8]. The previously published database known as the SASP Atlas, which used different genotoxic stressors in human fibroblasts and epithelial cells gave insight into these complexities [9]. This atlas shows that the core components associated with the soluble portion of the SASP are involved in altered extracellular matrix (ECM) organization, actin cytoskeleton and integrin interactions. Furthermore, in some fibrotic diseases, including pulmonary fibrosis, senescent cells and sSASP have been found to actively exacerbate fibrosis [10]. These results provide potential evidence that SASP factors remodel the ECM environment and modify the environment of neighboring cells.

Cell stiffness is sensed within tissues when cells exert force on ECM proteins via integrins, applying tension and meeting resistance that reflects the stiffness of the tissue [11]. Stiffer tissues in fibrosis cause cells to exert greater contractile forces, activating adhesion molecules, ion channels, cytoskeletal proteins, and associated signaling pathways to propagate mechanical stimuli through a process called mechanotransduction. The stiffness of young and healthy parenchymal tissues is on average 2 kPa [12]. However, in pathological conditions such as idiopathic pulmonary fibrosis, tissue rigidity elevates significantly, reaching as high as 50 kPa at advanced states [11,13].

Although mechanical forces certainly impact cellular function and phenotypes, little is known about how alteration of the physical ECM environment changes cellular senescence and the SASP. Our study investigates how mechanical tension, in the form of cell culture substrate rigidity, affects senescent cell makers and their soluble secretory phenotypes. We cultured IMR-90 human primary pulmonary fibroblasts on substrates mimicking physiological (2 kPa) and fibrotic (50 kPa) stiffness, as well as on conventional plastic plates (∼3 GPa) [14,15], followed by treatment with doxorubicin. We discovered unique secretory protein profiles and transcriptomic signatures that are modulated by high stiffness. Using a computational meta-analysis approach to mine published human interstitial lung disease single-cell RNA sequencing datasets, we confirmed that these genes are highly expressed in disease samples and strongly correlate with mechanotransduction and cellular senescence gene transcriptomic signatures. Thus, mechanical force shapes cell senescence and its soluble secretory products are linked to the pathogenesis of pulmonary fibrosis diseases.

## Results

### Cell substrate stiffness modulates senescence phenotypes in fibroblasts

We have previously optimized a cell culture system consisting of polydimethylsiloxane (PDMS) hydrogel polymerized on plastic tissue culture plates [16] with a fibronectin coating to mimic physiological substrate environments. The mechanical properties of hydrogels can be altered by changing the base: curing agent ratio, allowing it to be precisely tuned to mimic physiological tensions [17–20]. Because the normal lung tissue has a median physiological stiffness close to 2 kPa, we have utilized our previously developed 2 kPa hydrogel protocol, which has been validated by atomic force microscopy [16].

To understand how substrate stiffness impacts senescence phenotypes, we induced senescence in primary human lung fibroblast IMR-90 cells using 250nM doxorubicin (doxo) treatment for 24 hours and allowed 6 days for the phenotypes to develop on 2 kPa hydrogel or plastic (∼3 GPa) plates [21,22]. Induction of senescence in IMR-90 cells was confirmed using senescence-associated β-galactosidase (SA-β-Gal) activity (Fig. 1A) and 5-Ethynyl-2-deoxyuridine (EdU) incorporation (Fig.1B), previously recognized as standard measures of cellular senescence [23]. Unlike cells in a senescent state, untreated cells can either be actively proliferating or in a quiescent state, which is reversible and allows them to easily re-enter proliferation. Therefore, it was important to have both non-senescent controls that did not receive doxo treatment: proliferating cells that were grown for 4 days due to the prevention of over-confluence (proliferating control), and cells that were made quiescent by incubation in 0.2% serum media for 2 days prior to staining (quiescent control). Untreated proliferating cells, compared to quiescent cells, had a higher percentage of EdU incorporation and no induction of SA-β-Gal activity, as expected (Fig. 1A-B). Also as expected, doxo-treated senescent cells showed lower EdU incorporation than both control cells and an induction of SA-β-Gal activity in our model (Fig. 1A-B). Remarkably, doxo-induced senescent cells cultured on 2 kPa physiological hydrogel showed reduction in SA-β-Gal staining compared to those cultured on plastic, suggesting a role for substrate stiffness in senescence phenotype (Fig. 1A). This idea was further supported when we measured mRNA levels of *CDKN1a* (p21), a gene often highly expressed in senescent cells, and found an elevated trend driven by a hard matrix stiffness (Fig.1C). Together these results suggest that environmental stiffness can modulate cellular senescence markers.

**Figure 1.**
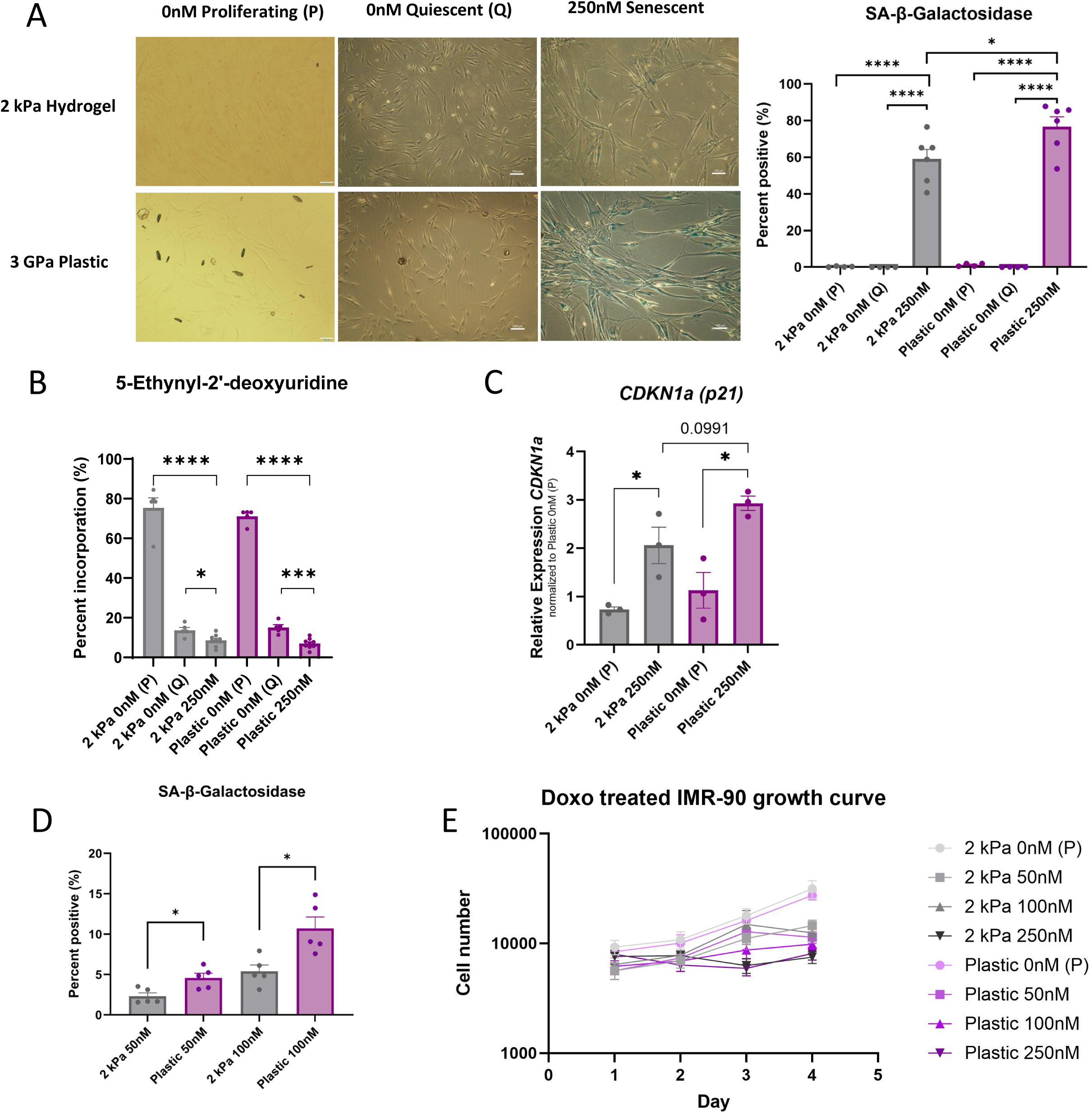
Substrate stiffness modulates senescence markers. A. Representative microscopy images showing IMR-90 human fibroblasts cultured on 2 kPa hydrogel and plastic (∼3 GPa) and stained with β-galactosidase (SA-β-gal) (left). SA-β-gal positive staining was quantified as a percentage of total cells (right). Senescence was induced using doxorubicin (250nM) for 24 hours. Senescent cells were imaged 6 days after treatment. Control proliferating cells (P) were untreated and grown for 4 days prior to staining. Control quiescent cells (Q) were also untreated but grown in low serum for 2 days to induce growth arrest prior to staining. Senescent cells were stained blue, and non-senescent cells negative cells had no color. (Scale bar = 100µm, n = 4 per untreated control and n=6 per doxo-treated condition). B. Quantification of 5-Ethynyl-2-deoxyuridine (EdU) staining of senescent and control IMR-90 fibroblasts cultured on different stiffnesses (n=5 per untreated control and n=9 per doxo-treated condition). EdU positive cells were quantified as a percentage of total cells. C. Quantitative PCR analysis of *CDKN1a (p21)* RNA levels in IMR-90 fibroblasts cultured on hydrogel and plastic. All values were normalized to plastic untreated proliferating (P) control cells (n=3 per condition). D. Quantification of SA-β-Gal activity in IMR-90 fibroblasts treated with 50nM and 100nM doxorubicin cultured on 2 kPa hydrogel and plastic (n=5 per condition). E. Growth curve of IMR-90 fibroblasts over four days, treated with increasing doses of doxorubicin cultured on 2 kPa hydrogel or plastic. Proliferating cells were used as untreated, non-senescent controls. No significance was observed at any day between comparable doses of doxorubicin in 2 kPa vs plastic (n=4 per group). Error bars represent SEM of biological variation, statistics evaluated via unpaired student t-test, dots represent biological replicates. (****p <0.0001, ***p <0.001, *p < 0.05).

To test whether stiffness modulated senescence induction occurs at lower levels of genotoxic stress, we next quantified SA-β-Gal activity in cells treated with lower doses of doxo of 50nM and 100nM. Under both doses, cells grown on plastic (∼3 GPa) showed significantly higher SA-β-Gal activity than cells cultured on the 2 kPa hydrogel (Fig.1D, Fig. S1). We then checked if the difference in SA-β-Gal level was due to a slower growth rate of cells cultured on the hydrogel, so cells were treated with 0nM, 50nM, 100nM and 250nM of doxo and tracked for cell growth (Fig.1E). We observed no difference in cell number between cells cultured on either stiffness level at any dose over the examined duration. IMR-90 cells treated with 250nM doxo did not proliferate, whereas the number of untreated cells more than tripled in 4 days (Fig. 1E). This suggests that induction of SA-β-Gal activity is likely not a consequence of stiffness-mediated changes to growth rate.

### Substrate stiffness modulates senescent cell associated secretory profile

To interrogate the composition of SASP of cells grown at different stiffnesses, we cultured 250nM doxo-induced IMR-90 cells for 8-10 days, allowing the production of senescent-associated secretory phenotypes (SASP)s in 2 kPa hydrogel and plastic (∼3 GPa) environments. Subsequently, cells were changed into serum-free media for 24 hours and the conditioned media containing soluble secretory proteins was collected. We performed label-free data-independent acquisition (DIA) mass spectrometry (MS) on collected soluble proteins, which is advantageous due to its sensitivity, accurate quantification, and large protein coverage by integrating the MS2 fragment ion chromatograms [24–26]. Senescent cells and their secretome are known to be heterogeneous [27,28]. Even *in vitro* cells that were uniformly induced for senescence can express substantially different genes across timepoints of development, as well as enter different cell cycle arrests, leading to varying levels of cytokine secretion [29,30]. For this reason, to enhance the robustness and reproducibility of detected proteins, we performed four independent MS experiments and each experiment included 4-6 biological replicates. The initial experiment was acquired on a TripleTOF 6600 mass spectrometer, and the subsequent three experiments on an Orbitrap Eclipse Tribrid platform.

Each proteomic experiment was individually analyzed (Fig. S2; Supplementary Tables 1-4) and then combined using the Fisher’s method on p-value results from four experiments to generate combined fold changes from the means (MetaFC) and adjusted P value [31] (Supplementary Table 5). In the combined result, proteins that were consistently and repeatedly identified were highly ranked, whereas proteins that were identified in only one of the experiments were ignored. We observed 1016 robustly differentially secreted proteins between plastic (∼3 GPa) vs. 2 kPa hydrogel cultured senescent cells, meeting the criteria of AdjP < 0.05, and MetaFC > 0.5 cutoff (Fig. 2A). Across all four experiments, the most significantly upregulated proteins under high stiffness conditions were Plectin (PLEC), collagen type VI alpha 3 chain (COL6A3) and Filamin family proteins, FLNA, FLNB, FLNC. Other cytoskeleton proteins like microtubule-associated proteins MAP4, MAP1B and MAP1A, talin (TLN1), lamin (LMNA) and spectrins, SPTAN1 and SPTBN1, and matrix proteins in the collagen family (COL) and laminins (LAMA) were also highly upregulated by the high stiffness of plastic. Interestingly, the softer physiological stiffness of 2 kPa upregulated the production of several inflammatory mediators like IL-6 and S100A8, extracellular matrix component TNC, and cytoskeleton organizers WDR1 and SEPTIN2, as well as the antioxidant enzyme superoxide dismutase SOD3, compared to plastic, suggesting a moderate level of unique stress response (Fig. 2A).

**Figure 2.**
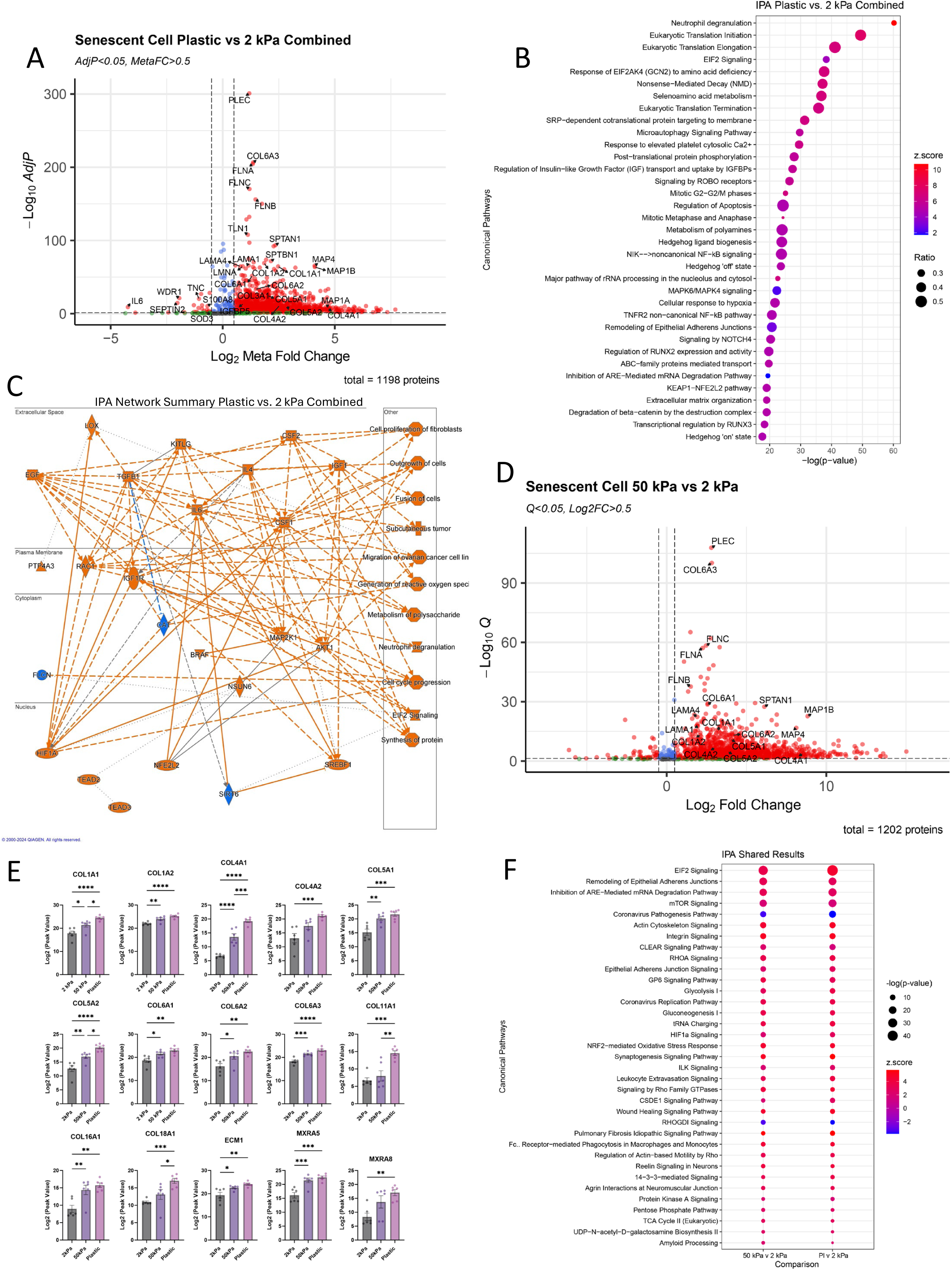
Stiffness drives differential secretion of senescence-associated secretory phenotype (SASP) proteins, measured by mass spectrometry (MS). A. Volcano plot showing proteins secreted into the culture supernatant by senescent IMR-90 fibroblasts treated with doxorubicin (250nM) cultured on plastic versus 2 kPa PDMS hydrogel as measured by MS. Four independent MS experiments were integrated to obtain this compiled secretory profile. Each MS experiment contained 4-6 biological replicates. B. Top 35 upregulated pathways in the secretome of senescent cells treated with doxorubicin (250nM) cultured on plastic compared to 2 kPa hydrogel generated using Ingenuity pathway analysis (IPA) software. C. Pathways and molecules enriched in the secretome of senescent cells treated with doxorubicin (250nM) cultured on plastic compared to 2 kPa hydrogel by IPA network analysis. D. Volcano plot showing proteins secreted into the culture supernatant by senescent IMR-90 fibroblasts treated with doxorubicin (250nM) cultured on 50 kPa vs. 2 kPa PDMS hydrogel as measured by MS (n=6 per group). E. Level of indicated collagen proteins and matrix-associated proteins secreted by 250nM doxorubicin-induced senescent fibroblasts cultured on different stiffness conditions, measured by MS (n=6). Error bars represent SEM of biological variation, statistics evaluated via one-way ANOVA, dots represent biological replicates. F. Overlapping pathways in the secretome of senescent fibroblasts treated with doxorubicin (250nM) when cultured on higher stiffness, evaluated using IPA software. IPA analysis cutoffs were set to p<0.05 and z-score >|1.5|.

To understand whether there were core pathways modulated by high stiffness, we performed pathway analyses on significantly altered secreted proteins using the Ingenuity Pathway Analysis (IPA) application. The most induced pathways by stiffness were neutrophil degranulation, many eukaryotic translation related processes, nonsense-mediated mRNA decay, and microautophagy signaling pathway (Fig. 2B). Although IL-6 protein was detected as upregulated by the 2 kPa hydrogel in MS, IPA predicted an upregulation of cytokines including IL-6, IL-4, TGFB1, KITLG, EGF and CSF2, due to other proteins in the same pathways being secreted under the high stiffness condition (Fig. 2C). Likewise, intracellular predictions by IPA highlighted the possible importance of RAC1, IGF1R and activation of AKT1, MAP2K1, and NSUN6 molecules, ultimately triggering the activation of master transcription factors HIF1A and NFE2L2 (Fig. 2B-C). These transcription factors are known regulators involved in mechanotransduction [32,33]. Additionally, well-known mechanotransducers, like TEAD2 and TEAD3, were also predicted to be activated by IPA analysis.

In pulmonary fibrosis, the local stiffness of the lung can increase up to 50 kPa [13]. To further investigate the stiffness modulation of senescent secretory profiles due to increases in stiffness in a disease-relevant condition, we also cultured IMR-90 cells on a 50 kPa PDMS hydrogel and induced senescence using 250nM doxo, and the soluble proteins were collected for MS analysis. We compared the secretome of senescent cells cultured on 50 kPa to those cultured on 2 kPa. A total of 1080 proteins met the differential cutoff (q-value <0.05, log2FC >0.5) (Fig. 2D). Proteins such as PLEC, COLs, LAMAs, SPTAN1, MAP4, and MAP1B, which were upregulated in plastic culture conditions, were also induced in the 50 kPa condition. The levels of many collagens (COL), particularly age-associated COL1 and COL6 [34], extracellular matrix protein 1 (ECM1), and matrix-remodeling-associated proteins (MXRA) showed a remarkably stiffness dependent increase across a range of stiffness from 2 kPa, 50 kPa and plastic (∼3 GPa) (Fig. 2E). To explore the biological effect of this differential secretion, we performed IPA analysis on the differentially secreted proteins between 50 kPa vs 2 kPa senescent cells and found many enriched pathways that were similar to the plastic vs 2 kPa profile (Fig. 2F).

Signaling molecules like mTOR, actin cytoskeleton, integrin, oxidative stress, and idiopathic pulmonary fibrosis pathways were upregulated in both comparisons, demonstrating that disease pathological stiffness (50 kPa) induced many similar SASP effects as supraphysiological plastic stiffness (Fig. 2F).

### Stiffness-modulated senescent secretory profile is influenced by control conditions

Performed alongside the 250nM doxo treated MS experiments, we also compared the secretory profiles of senescent cells with two different types of non-senescent controls: proliferating cells and quiescent cells. The goal was to identify the secretory profile unique to the senescence state under differential stiffness conditions, compared to quiescent and proliferation states.

Consistent with the senescent cell culture, we seeded these control cells either on plastic plates or 2 kPa hydrogel and collected their secretory proteins for MS analysis. Upon comparing plastic-cultured and 2 kPa hydrogel-cultured senescent cells with their respective proliferating control cells, we observed differential secretion of proteins such as the beta-galactoside-binding protein Galectin 7 (LGALS7), recycling endosomal phosphatase that facilitates YAP activation (PPP1R12A), immunophilin protein targeted by rapamycin (FKBP3), MAP1B, and nestin (NES) (Fig. 3A). Senescent cells on high stiffness induced a number of proteins, among which many were downregulated on softer matrix. However, 2 kPa substrate also exclusively upregulated a few proteins like Glypican-6 (GPC6) and mannose-associated serine protease 1 (MASP1). On the other hand, many proteins were also upregulated in senescent cells regardless of stiffness conditions, and these proteins included previously documented sSASP proteins, TIMP1 [9], secreted Frizzled-related protein 1 (SFRP1) [35,36], Tubulin beta-2A chain (TUBB2A) [37], extracellular matrix protein 1 (ECM1) [38] and CD109 [39] (Fig. 3B). In total, we detected 32 proteins that were consistently upregulated in senescent cells compared to proliferating cells under both stiffness conditions. This finding suggests a profile of stiffness-induced soluble SASPs that were also moderately modulated by doxo. Environmental stiffness resulted in varying sSASP profiles when doxo-induced senescent cells were compared to proliferating controls.

**Figure 3.**
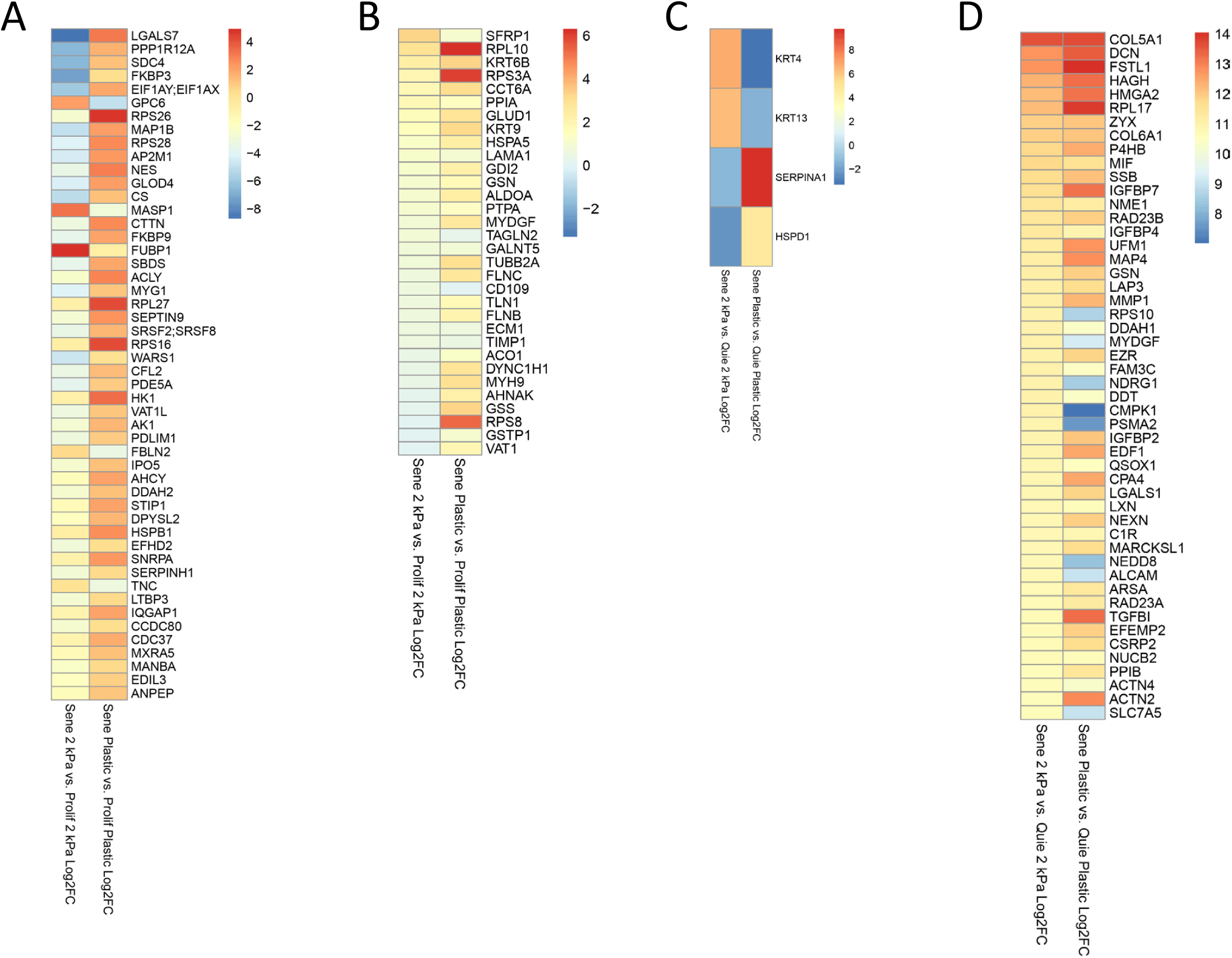
Differential stiffness induced senescent secretory profiles compared to proliferating and quiescent untreated control IMR-90 fibroblasts measured by MS. Heatmaps indicating the log fold change of secreted proteins from senescent cells compared to those from corresponding non-senescent controls. A. The top 50 differentially secreted proteins in senescent IMR-90 fibroblasts driven by stiffnesses compared to untreated proliferating controls. Red indicates upregulated secretion in senescent cells. Blue indicates upregulated secretion in proliferating untreated control cells. B. Shared upregulated proteins produced by senescent cells cultured on plastic or 2 kPa compared to untreated proliferating controls. All of the listed proteins had positive log fold change (logFC). C. Differentially secreted proteins of senescent cells cultured at plastic or 2 kPa compared to corresponding quiescent untreated controls. D. Top 50 upregulated proteins produced by senescent cells ranked by 2 kPa hydrogel culture condition compared to quiescent untreated controls. Both red and blue indicate positive log fold change (logFC).

We proceeded to compare senescent cells with quiescent controls cultured on plastic, as well as the corresponding groups cultured on 2 kPa hydrogel. Through this comparison, we detected a degree of stiffness-modulating effect. Serpin family A member 1 (SERPINA1) and heat shock protein family D member 1 (HSPD1) were upregulated by senescent cells only on plastic, whereas keratin 13 and 4 (KRT13, KRT4) were upregulated by senescent cells only on soft 2 kPa gel (Fig. 3C). However, the significance of these keratin protein changes is unknown as they are also common contaminates from sample handling. Senescent cells cultured on either stiffness, compared to quiescent controls, secreted many known sSASPs including IGFBP families, COL families and high mobility group AT-hook (HMGA1,2) [40] (Fig. 3D). While stiffness did not impact so much the type of protein produced in senescent vs. quiescent cells, it did impact the level of upregulation of many sSASP proteins, demonstrating the differential expression capacity of sSASPs modulated by various stiffness conditions.

### Transcriptomic analysis reveals a stiffness driven gene expression profile

To better understand the mechanisms by which matrix stiffness impacts senescence secretory profiles, we proceeded to investigate the effects of various matrix stiffness levels on transcription. We conducted bulk mRNA sequencing on 250nM doxo-induced IMR-90 senescent cells cultured on either 2 kPa PDMS hydrogels or ∼3 GPa plastic plates. Senescent cells cultured on plastic exhibited upregulation of 17 genes and downregulation of 7 genes, with a cutoff of adj.p < 0.05 and log2FC > 0.5 compared to those cultured on 2 kPa hydrogel (Fig. 4A). Proteins like COL1A1, COL3A1 and IGFBP5 that were induced in the MS results (Fig. 2A) were also observed as upregulated by mRNA transcription at these statistical cutoffs (Fig. 4A). To compare the level of similarity between proteomic and transcriptomic results, we repeated IPA pathway analysis on mRNA data using differential expressed genes (DEG) with a raw p < 0.01, due to the limited number of genes at the adj.p <0.05 cutoff. High stiffness drove collagen biosynthesis, extracellular matrix organization, syndecan interaction, platelet-derived growth factor (PDGF) and glycoprotein VI platelet (GP6) signaling, and fibrosis signaling pathways (Fig. 4B). The high rigidity of plastic consistently induced multiple extracellular matrix (ECM)-related terms in both proteomic and transcriptomic analyses (Fig. S3A). On the other hand, 2 kPa hydrogel showed a strong induction in cholesterol biosynthesis processes at a transcript level, as well as other metabolic processes (Fig. 4B). Similarly to proteomic studies, the IPA analysis using DEGs predicted the activation of direct mechanosensitive molecules, this time inducing YAP1 and MRTFA pathways under plastic stiffness (Fig. 4C). This suggests that mechanotransduction pathways may play a pivotal role in governing changes to specific sSASP products under varying stiffness conditions.

**Figure 4.**
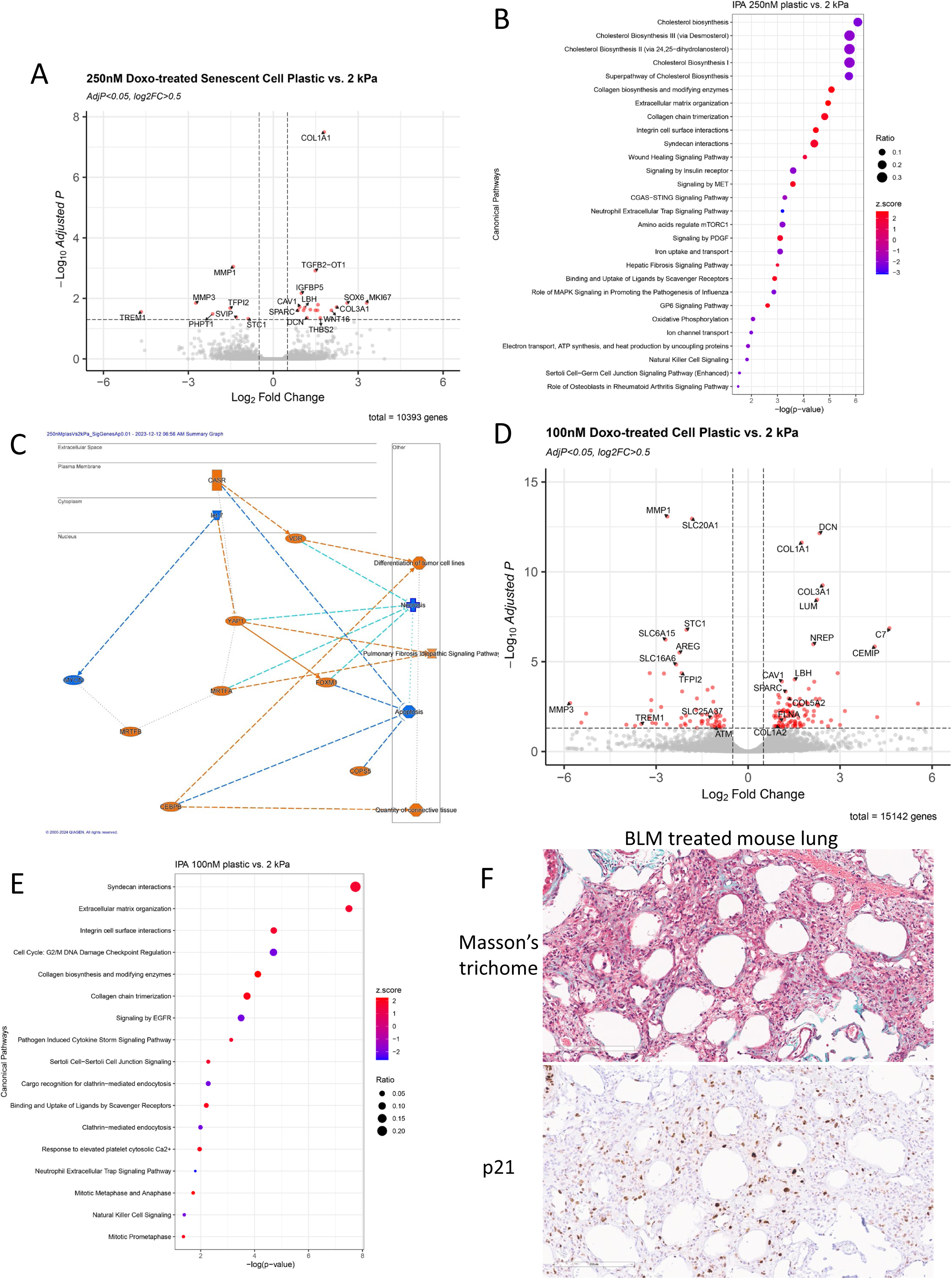
RNA sequencing analysis shows differential senescence gene transcription under different stiffness conditions. A. Volcano plot of differentially expressed genes (DEGs) in senescent IMR-90 fibroblasts induced using doxorubicin (250nM) and cultured on ∼3 GPa plastic versus on 2 kPa hydrogel (n=3 per condition). B. Ingenuity pathway analysis (IPA) of DEGs comparing senescent cells treated with 250nM doxorubicin cultured on plastic versus on 2 kPa hydrogel. IPA analysis cutoffs were set to p<0.05 and z-score >|1.5|. C. Differential pathways and molecules enriched in DEGs from doxorubicin (250nM) treated senescent cells cultured on plastic compared to 2 kPa hydrogel generated by IPA network analysis. D. Volcano plot of DEGs in fibroblasts treated with 100nM doxorubicin cultured on plastic vs 2 kPa hydrogel (n=3 per condition). E. Pathway analysis (IPA) of DEGs in fibroblasts treated with 100nM doxorubicin cultured on plastic vs 2 kPa hydrogel. IPA analysis cutoffs were set to p<0.05 and z-score >|1.5|. F. Immunohistochemistry images of lungs from bleomycin-treated mice stained with Masson’s trichome (top) and p21 (bottom). Fibrosis is shown in blue, including blue streaks or blue wisps (top), while p21 staining is in brown (bottom). Scale bar = 200µm.

Previously, we saw significant differences in SA-β-Gal staining at 50nM and 100nM doxo treatment between plastic and 2 kPa hydrogel conditions (Fig. 1D). In order to explore the potential subtle mechanisms that regulate senescence markers in different stiffness conditions, we chose the 100nM doxo dose for its partial induction of senescent markers and performed bulk mRNA-sequencing on 100nM doxo treated cells cultured on plastic or 2 kPa hydrogel (Fig. 4D). Even using this lower dose of doxo, multiple collagen genes and genes known to be involved in collagen fibrillogenesis [41] like *DCN* and *LUM*, exhibited robust upregulation in response to hard substrate (Fig. 4D). Conversely, *MMP*s and several solute carrier family members (*SLC*) were induced by the physiological 2 kPa matrix. Pathway analysis using DEGs revealed that the stiff plastic substrate activated pathways related to ECM organization, collagen and integrin interaction (Fig. 4E), which was consistent with the results observed in cells treated with the higher 250nM doxo dose (Fig. 4B). However, a unique term of Cell cycle: G2/M damage checkpoint regulation, driven by ataxia telangiectasia mutated (*ATM*) activity, was strongly induced by 2 kPa hydrogel (Fig. S3B). This suggests that cells cultured at physiological stiffness may activate unique compensatory mechanisms in response to genotoxic agents or enter senescence through distinct pathways compared to cells under higher supraphysiological stiffnesses.

### Bleomycin-induced fibrosis mouse lungs showed clusters of p21 positive cells in areas of fibrosis

Given the enrichment of collagen and fibrosis-related proteins and genes produced by senescent cells in a higher stiffness environment, we next sought to see if there was evidence of senescent cells colocalizing to regions of fibrosis *in vivo* during senescence-associated disease. To assess this idea, we utilized a bleomycin-induced pulmonary fibrosis model of idiopathic pulmonary fibrosis. Mice received one dose of bleomycin injection, and the lung tissues were imaged with p21 and Masson’s trichrome stains (Fig. S4A). The control mice showed little blue stain on trichrome, indicating little to no fibrosis (Fig. S4B). There were only a few p21 positive cells on the slide. In contrast, the bleomycin-treated mice showed regions of interstitial thickening with fibrosis and alveolar architectural distortion (Fig. 4F). Treated lungs had numerous p21 positive cells scattered around the tissue, consistent with senescence induction. P21 cells were also notably observed associated with regions of fibrosis, as well as regions of higher architecture distortion (Fig. 4F and Fig. S4C), suggesting the potential for *in vivo* relevance of stiffness associated SASPs.

### Upregulation of stiffness-associated sSASP genes in human lung diseases

We have identified distinct secretory profiles of senescent fibroblasts within higher stiffness environments and the presence of p21+ cells associated with fibrosis in mouse models of pulmonary fibrosis. Consequently, we were intrigued by the possibility that these identified stiffness-associated proteins and genes may be linked to human diseases governed by increased stiffness and cell senescence within the lung. Such increases in stiffness and senescence are often seen in pulmonary fibrosis diseases, including idiopathic pulmonary fibrosis, potentially playing a role in the pathogenesis [10,42]. To explore this hypothesis, we reanalyzed 4 previously published single-cell RNA-sequencing datasets related to lung fibrosis diseases (Fig. S5A), encompassing a total of 66 human lung samples using integrative Seurat analysis [43–48]. The combined dataset consists of 29 healthy controls and 47 cases of interstitial lung disease (ILD) that are predominantly idiopathic pulmonary fibrosis cases, but also include other ILDs like systemic sclerosis, chronic hypersensitivity pneumonitis and sarcoidosis.

The dimensionality reduction method uniform manifold approximation and projection (UMAP) of the integrated data identified 31 cell types (Fig. S5B), employing the annotation method detailed in one of the datasets [46]. Based on this annotation, the fibroblast population was further classified into four subtypes: myofibroblast, fibroblast, PLIN2+ fibroblast and HAS1 high fibroblast (Fig. S5C). Given our specific interest in fibroblasts, we incorporated transcripts from all four subtypes in the subsequent analysis. We then mapped the gene expression patterns of control and ILD samples against the top-secreted protein profiles identified in our MS results (data from Fig. 2). Specifically, we selected the top 30 upregulated proteins from senescent cells treated with 250nM doxo and cultured on plastic compared to those on 2 kPa hydrogel.

Additionally, we selected the top 30 upregulated proteins between cells that were treated with 250nM doxo and cultured on 50 kPa versus 2 kPa hydrogel. Notably, ILD led to a significant enrichment in these stiffness-driven sSASP genes, despite the stark difference of using doxo-induced *in vitro* senescence secretome as model (Fig. 5A). Subsequently, we focused on the upregulated genes (adjp < 0.1), totaling 22 genes, derived from our transcriptomic results of senescent cells cultured on plastic compared to those on 2 kPa gel (data from Fig. 4A), the ‘hard’ DEGs. We compared the average gene expression level of the listed gene in both control and ILD samples, revealing an enrichment in plastic induced DEGs in disease compared to healthy controls (Fig. 5B). We also analyzed the enrichment of genes differentially expressed in 2 kPa senescent cells, namely ‘soft’ DEGs, and observed significant enrichment though to a lesser extent than plastic induced ‘hard’ DEGs (Fig. S6, and Fig. 5B). However, this enrichment in “soft” DEGs in ILD is likely due to the enrichment of overall SASP genes, such as *MMP*s.

**Figure 5.**
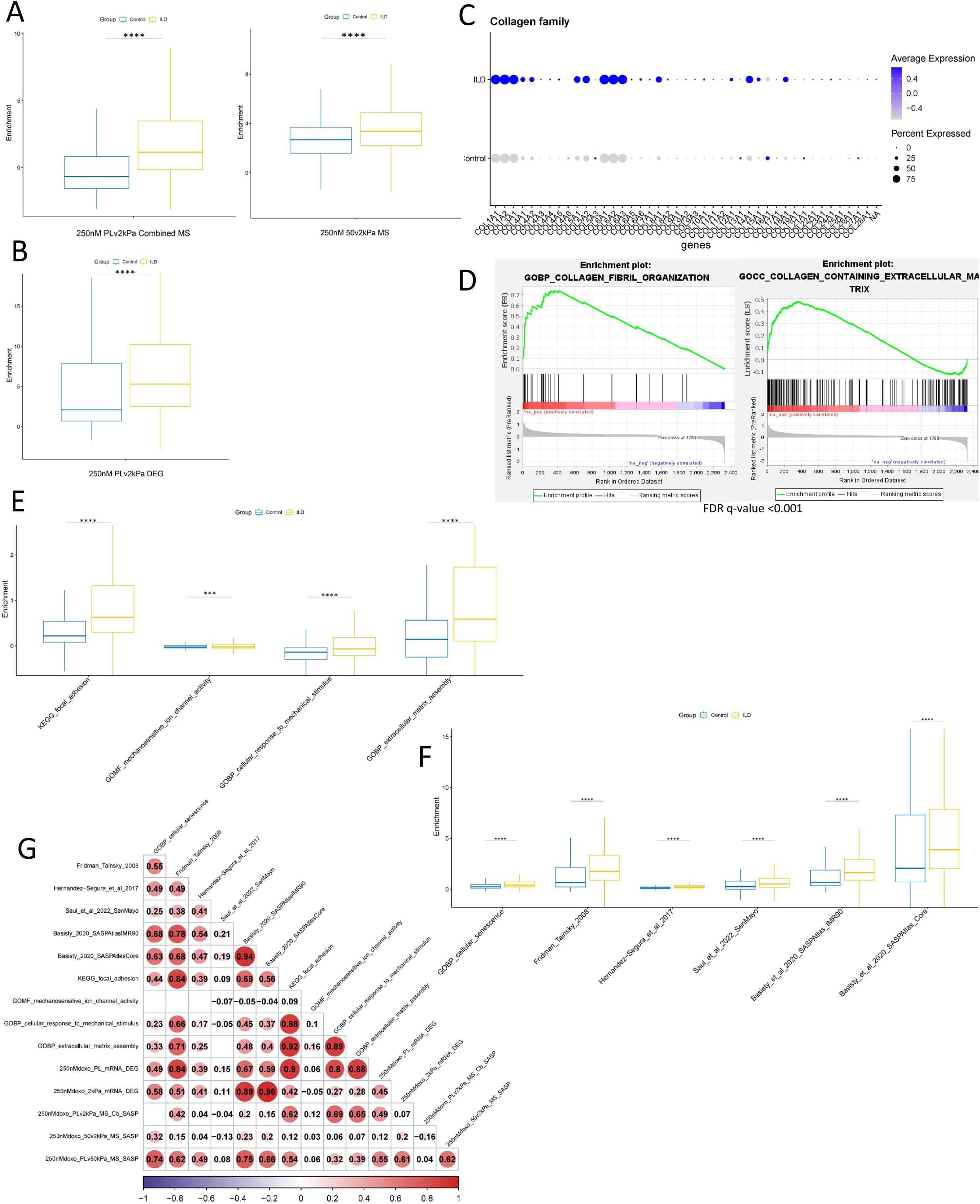
Comparative meta-analysis using four single-cell RNA sequencing datasets from human lung diseases shows induction of stiffness-modulated proteins and genes, senescent genes, and mechanotransduction genes. Statistics calculated using the Wilcoxon Rank Sum test (****p <0.0001). A. Enrichment scores of interstitial lung disease (ILD) and healthy control fibroblasts in plastic versus 2 kPa and 50 kPa versus 2kPa using secretory proteomic profiles generated from mass spectrometry. B. Enrichment score of interstitial lung disease (ILD) and healthy control fibroblasts in plastic versus 2 kPa DEG using transcriptional signatures generated from RNA-sequencing analysis. C. Transcriptional changes in multiple collagen genes as indicated, comparing ILD fibroblasts to healthy control. D. Enrichment of collagen-related terms in ILD fibroblasts by gene set enrichment analysis (GSEA). E. Enrichment scores of mechanotransduction and ECM-related genes and pathway terms in ILD and healthy control fibroblasts. F. Enrichment scores of ILD and control fibroblasts in senescence related genes and pathway terms, with relevant study data set indicated. G. Pearson correlation co-efficient mapping comparing human fibroblast enrichment scores in senescence genes, mechanotransduction related genes, stiffness-driven protein and gene signatures across gene sets as indicated.

Because collagens were robustly upregulated in both stiffness-related sSASPs and DEGs, we examined the expression of 42 collagen proteins in the fibroblast transcriptome of ILD vs. control samples. The ILD cells exhibited elevated expression of several collagen genes, with notable increases in *COL1A1, COL1A2, COL6A1, COL6A2, and COL6A3* (Fig. 5C). Further supporting this finding, we performed gene set enrichment analysis (GSEA) and confirmed that collagen-related gene ontology (GO) terms were significantly enriched exclusively in ILD fibroblasts (Fig. 5D).

Mechanotransduction is a process by which physical stimuli are propagated into the cell through mechanosensitive molecules. Therefore, we suspected the involvement of mechanotransduction pathways in stiffness modulated sSASP profile. To test this hypothesis, we mapped the gene expression of control and diseased fibroblasts against well-annotated mechanotransduction and extracellular matrix gene sets from Molecular Signatures Database (details of gene sets selected from literature were listed in Supplementary Table 6) [49–51]. ILD samples exhibited stronger enrichment compared to healthy controls for GO and KEGG terms such as focal adhesion, cellular response to mechanical stimulus, extracellular matrix assembly and mechanosensitive ion channel (Fig. 5E). Although Hippo signaling YAP and TAZ molecules are well-known mechanosensors, their nuclear translocation is more relevant to mechanical stimuli than their transcription [52]. Additionally, Hippo signaling gene sets include many YAP/TAZ upstream and downstream genes, which are less mechanosensitive. For these reasons, the Hippo pathway gene set was not used in our analysis. ILD fibroblasts also showed strong enrichment for many highly cited senescence gene signatures [9,53–55], as previously described [10] (Fig. 5F).

Next, to explore potential expression correlation in fibroblasts between gene sets associated with cellular senescence and those related to mechanotransduction, including those from our *in vitro* model, we computed the Pearson correlation between these gene sets (Fig. 5G).

Senescence and mechanotransduction related gene sets showed a high level of intra-correlation, attributed to shared biological pathways and a few overlapping genes (Fig. S7), with the exception of SenMayo [54] and mechanosensitive ion channel. We found that SenMayo showed poor correlation with other senescent gene sets, and had a weak correlation with all selected mechanotransduction associated gene sets. Mechanosensitive ion channels also exhibited a weak correlation with other gene sets, possibly because their transcript levels are less influenced by senescence or mechanical stimuli; rather, it is anticipated that ion channels’ activation at the protein level will be more affected. Interestingly, there is an overall positive correlation between selected mechanotransduction related gene sets and senescence gene sets. The gene signature from Fridman *et al.* [53] demonstrated a notably strong positive correlation with mechanotransduction-related gene sets. Similarly, Basisty *et al.*’s [9] protein signatures exhibited a correlation of > 0.4 with mechanotransduction related gene sets. This dataset showed a trend of co-expression between these senescence and mechanotransduction signatures. Last, we tested the correlation of stiffness driven top sSASPs and DEGs used in our manuscript to the senescence and mechanotransduction associated gene sets. Stiffness driven signatures were positively correlated with both groups of genes. The plastic vs. 2 kPa sSASPs were highly correlated with mechano-related genes with values > 0.6. The ‘hard’ DEG from 250nM doxo plastic vs. 2 kPa was highly positively correlated with almost all senescent gene sets and mechano-related gene sets. On the other hand, the 2kPa gel ‘soft’ DEG compared to the plastic DEG showed reduced levels of correlation on all mechanotransduction-related gene sets. These findings again support the notion that mechanotransduction pathways are correlated with, and may potentially alter, sSASP outputs.

## Discussion

Here we demonstrate that substrate stiffness impacts senescent-associated secretory phenotypes (SASP)s, gene expression and senescence-associated β-galactosidase activity in the primary lung fibroblast cell line IMR-90. We also show that high stiffness induced soluble SASP and gene profiles from our model are highly expressed in human interstitial lung diseases, and they have strong correlation to existing cellular senescence and mechanotransduction gene sets. These results provide evidence that cellular senescence and SASPs are sensitive to different mechanical environments, and they may have potential interactions in diseases where tissue stiffness is altered. Consistently, we observed p21+ cells present in regions of fibrosis and lung architectural distortion in mouse model of idiopathic pulmonary fibrosis.

Mechanical forces serve as crucial signals for cells. Our data demonstrates that senescent cells cultured on fibrosis-mimicking stiffness level (50 kPa), as well as supraphysiological plastic (∼3 GPa), secrete significantly more ECM proteins and exhibit a robust secretory profile compared to those on 2 kPa stiffness, underscoring the sensitivity of cells to their mechanical environment. One caveat explanation is that higher stiffness may induce more senescent cells, as seen with the SA-β-gal assay on plastic vs. 2kPa, thereby partially contributing to the upregulated SASP secretion. However, our core signatures still correlated across human senescence and mechanotransduction profiles, suggesting that core features of our stiffness SASP are at play in human age-related fibrotic disease. Indeed, throughout the aging process, various tissues and organs undergo changes in stiffness, coinciding with the accumulation of senescent cells. For instance, the elastic modulus of lung parenchymal tissue can nearly double with age, even in the absence of disease [12]. Our research and others have shown that stiffness changes can reprogram immune cell metabolism and alter inflammatory output [16,56,57]. Despite the unclear mechanisms underlying their pathologies, many lung fibrosis diseases exhibit a positive correlation with age. Our data suggests that environment stiffness may work with dominant age-related stressors, like genotoxic stress, to facilitate cell senescence; cell senescence then contributes to SASP production shaped by environmental force, leading to extracellular remodeling, favoring collagen deposition and fibrosis when stiffness increases.

Consistently, our data showed a reproducible and significant upregulation of collagen family genes, both at the transcript and protein levels, in response to high stiffness. Collagens are highly involved in fibrosis as an excess in their synthesis relative to their degradation leads to their accumulation, which promotes fibrosis. While studies have reported differential expression of collagen by senescent cells, evidence suggests that collagens may also modulate cellular senescence [58,59]. Furthermore, multiple collagens including COL1A1, COL6A1 and COL6A2 exhibit a strong positive association with age in the human lung [34]. Collectively, these findings suggest that the high stiffness-induced collagen expression and secretion observed in our study likely play a significant role in the interplay between senescence and fibrotic diseases.

Recent research has confirmed the upregulation of collagen in response to high stiffness while also revealed differential expressions patterns of cyclin-dependent kinase inhibitors (CDKN) and cytokines in cells cultured on hydrogels of varying stiffness. A study using primary human lung fibroblasts on a polyacrylamide hydrogel with stiffness of 1 kPa mimicking healthy and 23 kPa mimicking fibrotic lungs, found that HDAC3 cytosol localization triggered upregulation of *COL1A1* and *CDKN1a* (*p21WAF1/CIP1)* in the fibrotic stiffness [60]. One study assessed replicative senescence of WI-38 human lung fibroblasts cultured on 0.5, 2, 16, 32 kPa PDMS gel and 2.3 GPa plastic at 4 weeks and observed the lowest level of SA-β-Gal on plastic and highest level of SA-β-Gal on 0.5kPa [61]. However, soft matrix lowered the mRNA expression of cytokine gene *IL-1* and had no impact on *CDKN1a*, *CDKN2a (p16INK4a)*, and *CDKN2b (p15INK4b).* Another study on lung fibroblasts used gelatin methacrylate substrates with stiffnesses of 5 kPa and 15 kPa, observing no changes in *CDKN1a* or *CDKN2a* gene expression, but an increase in *COL1A1* with increasing stiffness [62], consistent with our observed increased collagen SASP under higher stiffness. Other studies that used fibroblasts from other origins have reported varying levels of SA-β-Gal, CDKNs and inflammatory cytokines when cultured on soft versus stiff substrates [63]. Because the materials and stiffnesses assessed in these studies were inconsistent, it is difficult to make direct comparisons. Moreover, variation in results may also point to differential impacts of mechanical forces on different inducers of senescence, such as replicative versus genotoxic. However, despite these variations, all studies have observed a general mechanosensitive nature of these markers, and when coupled with our findings, pose a novel potential confounding factor in senescence identification.

The conventional view of senescence portrays it as an irreversible cell cycle arrest. Yet, ongoing research suggests that senescence is a complex stress response. Mechanisms that induce apoptosis can also drive cellular senescence [64]. ATM kinase is an important protein in DNA damage response that also plays an important role in fine tuning the balance between senescence and apoptosis. We found that ATM is upregulated by cells cultured on 2 kPa hydrogel treated with 100nM doxo, which drove the positive enrichment for cell cycle checkpoint regulation, even though a lower level of SA-β-gal activity is observed. Although ATM kinase is generally thought to promote senescence, studies have found that activation of ATM kinase activity can alleviate replicative senescence [65,66]. Though we identified ATM enriched in softer physiological stiffness gels, ATM kinase has also been shown to be modulated by mechanical stretching [67], and YAP/TAZ signaling, facilitating senescence [68]. Thus, more work is needed to better understand the role of ATM in senescence, as well as the precise mechanism it uses to interact with mechanotransduction pathways.

In many SASP studies, the choice of control conditions in comparison to senescence is often a subtle detail that is overlooked. In our study, we investigated two controls, revealing that using either proliferating or quiescent cells significantly influence the results. When senescent cells were compared to proliferating cells, stiffness had a substantial impact on differential secretion. Conversely, senescent cells compared to quiescent cells exhibited a less pronounced differential secretion effect to stiffness, though the magnitude of secretion was still markedly influenced by stiffness. These findings underscore the importance of selecting appropriate controls in proteomic analysis.

A significant amount of research has been dedicated to accurately characterizing the SASP, including the development of the SASP Atlas aimed at identifying reliable biomarkers and therapeutic targets [9,69–71]. It is now understood that the SASP encompasses not only soluble factors, but also extracellular vesicles [72], which play a vital role. However, much of work in the field has been conducted using cells cultured under supraphysiological forces that deviate from *in vivo* biology. Given the challenges associated with measuring *in vivo* SASP, as well as its heterogeneous nature, it may be important that future iterations of the SASP Atlas can be better refined to mimic *in vivo* physiology with changes to culture techniques to simulate physiological forces on cells.

In summary, our findings suggest that senescence development in fibroblasts are responsive to the mechanical environment. As aging progresses, we hypothesize that cells experience a chronically altered stiffness environment due to tissue scarring or the natural aging processes. As senescent cells emerge with accumulating stresses in the lung, cells continue to perceive distinct mechanical cues, thereby triggering mechanotransduction mechanisms and inducing an altered secretory profile. These senescent cells contribute to the progression of fibrosis by secreting sSASPs that are enriched in ECM proteins. However, more work is needed to figure out the precise mechanosensitive mechanisms at play in the modulation of senescence and SASP. For instance, the extent to which the tissue aging mechanical environment, including exhaustion of mechanotransduction signals across time, contributes to disease manifestation remains unclear. In fact, the increasing identification of mechanosensors as effective therapeutic targets for age-related diseases further underscores their importance in the aging process [73–75]. It will also be important to better map out the effects of 3D viscoelasticity models on cell senescence and the sSASP, as such models may better mimic *in vivo* physiology than the 2D hydrogels used in the current study [76–78]. Furthermore, despite showing strong enrichment of high stiffness sSASP in human fibroblasts under fibrosis, we were limited to mapping protein signatures to existing single-cell mRNA databases rather than proteomic data. Therefore, future work is required to directly measure stiffness-induced SASP *in vivo* and establish links between stiffness-induced SASP and mechano-modulation of senescence, to better understand their involvement in fibrotic diseases and aging. Overall, our work provides evidence of mechanical force tuning of cell senescence and its secretory products, with potential relevance to age-related fibrotic disease.

## Methods

### PDMS Hydrogel-coated plates

The gels were prepared using Dow Corning Sylgard 527 silicone dielectric gel (Part A and B, Ellsworth Adhesives, Germantown, WI). Part A and part B components of the gel were mixed at the following ratios to obtain the appropriate tensions. For 2 kPa gel, the ratio of A:B was 1.2 and for 50 kPa gel the ratio of A:B was 0.3. The gel was poured onto polystyrene cell-culture plates (Nunc, Thermo Fisher Scientific, Waltham, MA; #140675) and incubated for3 24 hours at 60°C. After solidifying, gels were coated with fibronectin (1μg/mL in PBS, Thermo Fisher Scientific, Waltham, MA; #10010023) for 1 h at 37°C followed by two washes with PBS. The gel-coated plates were used immediately after washing. The stiffness of 2 kPa and 50 kPa gel was validated by atomic force microscopy [17].

### Human primary cell lines and cell culture

IMR-90 primary human lung fibroblasts (ATCC, Manassas, VA; #CCL186) were cultured in Dulbecco’s Modified Eagle Medium (Thermo Fisher Scientific, Waltham, MA; #12430054) supplemented with penicillin-streptomycin (5,000 U/mL; Thermo Fisher Scientific, Waltham, MA; #15070063) and 10% fetal bovine serum (Thermo Fisher Scientific, Waltham, MA; #2614079). Cells were seeded on either conventional polystyrene cell-culture treated 6-well plate (Nunc, Thermo Fisher Scientific, Waltham, MA; #140675), 2 kPa or 50 kPa PDMS-coated plates for overnight before any following treatment.

### Doxorubicin induced senescence induction

Cells were cultured in above mentioned media containing either 50, 100, or 250 μM doxorubicin, or vehicle (DMSO) for 24 hours. Subsequently, cells were cultured for at least 8 days to allow development of the senescent phenotype. Proliferating control cells were cultured in complete media for 4 days. Quiescence was induced in control cells by culturing in complete media for 1 day and 0.2% FBS serum media for the subsequent 2 days.

### Isolation of secreted soluble proteins

After the treatment period described above, cells were washed with PBS (Thermo Fisher Scientific, Waltham, MA; #10010023) and placed in serum- and phenol red-free DMEM (Thermo Fisher Scientific, Waltham, MA; #21063029). Conditioned media was collected after 24 hours.

Conditioned media was then centrifuged at 10,000*g* at 4°C for 30 minutes to remove debris and stored in -80°C for proteomic analysis.

### Proteomic analysis

#### Sample Concentration

6-8 mL of conditioned media (Experiment 1: n = 4 for each condition – one replicate was obtained by pooling the conditioned media from 3 wells) or 2 mL of conditioned media (Experiment 2: n = 6 for each condition; Experiment 3: n = 6 for each condition; Experiment 4: n = 3-4 for each condition; with for all three experiments, one replicate was obtained from a single well) from IMR-90 cultures under different tensions and doxo treatments were concentrated using 3 kDa molecular cut-off filters (Millipore Sigma, Burlington, MA) and protein content was quantified using the bicinchoninic acid assay (BCA; Thermo Fisher Scientific).

#### Protein Digestion and Desalting

Aliquots of concentrated secretome from each sample were reduced using 20 mM dithiothreitol in 50 mM triethylammonium bicarbonate buffer (TEAB) at 50°C for 10 min followed by 10 min at room temperature (RT), and alkylated using 40 mM iodoacetamide in 50 mM TEAB at RT in the dark for 30 min. Samples were acidified with 12% phosphoric acid to obtain a final concentration of 1.2% phosphoric acid. Seven volumes of S-Trap buffer consisting of 90% methanol in 100 mM TEAB at pH ∼7.1 were added and samples were loaded onto the S-Trap mini or micro spin columns. The entire sample volume was spun through the S-Trap micro spin columns at 4,000 x *g* and RT, binding the proteins to the micro spin columns. Subsequently, S-Trap micro spin columns were washed twice with S-Trap buffer at 4,000 x *g* at RT and placed into clean elution tubes. Samples were incubated for one hour at 47°C with a solution of sequencing grade trypsin (Promega, San Luis Obispo, CA) dissolved in 50 mM TEAB at a 1:25 (wt/wt) enzyme:protein ratio. Afterwards, trypsin solution was added again at the same ratio, and proteins were digested overnight at 37°C.

Peptides were sequentially eluted from mini or micro S-Trap spin columns with 50 mM TEAB, 0.5% formic acid (FA) in water, and 50% acetonitrile (ACN) in 0.5% FA. After centrifugal evaporation, samples were resuspended in 0.2% FA in water and desalted with Oasis 10 mg Sorbent Cartridges (Waters, Milford, MA). The desalted elutions were then subjected to an additional round of centrifugal evaporation and re-suspended in 0.2% FA in water at a final concentration of 1 µg/µL. Finally, indexed Retention Time standard peptides (iRT, Biognosys, Schlieren, Switzerland) [79] were spiked in the samples according to manufacturer’s instructions.

#### Mass Spectrometric Analysis of Experiment 1 (TripleTOF 6600 MS)

LC-MS/MS analyses were analyzed by reverse-phase HPLC-ESI-MS/MS using the Eksigent Ultra Plus nano-LC 2D HPLC system (Dublin, CA) combined with a cHiPLC system directly connected to an orthogonal quadrupole time-of-flight TripleTOF 6600 mass spectrometer (SCIEX, Redwood City, CA). Typically, mass resolution in precursor scans was approximately 45,000, and fragment ion resolution was approximately 15,000 in “high sensitivity” product ion scan mode. After injection, peptide mixtures were transferred onto a C18 pre-column chip (200 μm × 6 mm ChromXP C18-CL chip, 3 μm, 300 Å; SCIEX) and washed at 2 μL/min for 10 min with the loading solvent (0.1% FA in water) for desalting. Peptides were transferred to the 75 μm × 15 cm ChromXP C18-CL chip, 3 μm, 300 Å (SCIEX) and eluted at 300 nL/min with a 3-h gradient using aqueous and acetonitrile solvent buffers.

All samples were analyzed by data-independent acquisition (DIA), specifically using variable window DIA acquisitions [25] In these DIA acquisitions, 64 windows of variable width (5-90 m/z) were passed in incremental steps over the full mass range (m/z 400-1,250) with an overlap of 1 m/z. The cycle time of 3.2 seconds included a 250-ms precursor ion scan, followed by acquisition of 64 DIA MS/MS segments, each with a 45-ms accumulation time (Supplementary Table 7). The variable windows were determined according to the complexity of the typical MS1 ion current observed within a certain m/z range using a SCIEX “variable window calculator” algorithm (more narrow windows were chosen in “busy” m/z ranges, wide windows in m/z ranges with few eluting precursor ions) [26]. DIA tandem mass spectra produce complex MS/MS spectra, which are a composite of all the analytes within each selected Q1 m/z window.

#### Mass Spectrometric Analysis of Experiment 2, 3 and 4 (Orbitrap Eclipse Tribrid MS)

LC-MS/MS analyses were performed on a Dionex UltiMate 3000 system coupled to an Orbitrap Eclipse Tribrid mass spectrometer (both from Thermo Fisher Scientific, San Jose, CA). The solvent system consisted of 2% ACN, 0.1% FA in water (solvent A) and 98% ACN, 0.1% FA in water (solvent B). For Experiment 2, proteolytic peptides (800 ng) were loaded onto an Acclaim PepMap 100 C18 trap column (0.1 x 20 mm, 5 µm particle size; Thermo Fisher Scientific) for 5 min at 5 µL/min with 100% solvent A. Peptides were eluted on an Acclaim PepMap 100 C18 analytical column (75 µm x 50 cm, 3 µm particle size; Thermo Fisher Scientific) at 300 nL/min using the following gradient of solvent B: 2% for 5 min, linear from 2% to 20% in 125 min, linear from 20% to 32% in 40 min, up to 80% in 1 min, 80% for 9 min, and down to 2% in 1 min. The column was equilibrated with 2% of solvent B for 29 min, with a total gradient length of 210 min. For Experiment 3 and Experiment 4, proteolytic peptides (500 ng for Experiment 3 and 400 ng for Experiment 4) were loaded on Acclaim PepMap 100 C18 trap column (75 µm x 20 mm, 3 µm particle size; Thermo Fisher Scientific) for 10 min at 2 µL/min with 100% solvent A. Peptides were eluted on an Acclaim PepMap 100 C18 analytical column (75 µm x 50 cm, 3 µm particle size; Thermo Fisher Scientific) at 300 nL/min using the following gradient of solvent B: 2% for 10 min, linear from 2% to 20% in 125 min, linear from 20% to 32% in 40 min, up to 80% in 1 min, 80% for 9 min, and down to 2% in 1 min. The column was equilibrated with 2% of solvent B for 29 min, with a total gradient length of 215 min.

All samples were acquired in DIA mode. Full MS spectra were collected at 120,000 resolution (AGC target: 3e6 ions, maximum injection time: 60 ms, 350-1,650 m/z), and MS2 spectra at 30,000 resolution (AGC target: 3e6 ions, maximum injection time: Auto, NCE: 27, fixed first mass 200 m/z). The isolation scheme consisted in 26 variable windows covering the 350-1,650 m/z range with an overlap of 1 m/z (Supplementary Table 8) [80].

### DIA-MS Data Processing and Statistical Analysis

For Experiment 1, DIA data were processed in Spectronaut v14 (version 14.10.201222.47784; Biosgnosys, Schlieren, Switzerland) using the panhuman library that provides quantitative DIA assays for 10,316 human proteins [81] and supplemented with scrambled decoys (library size fraction of 0.1). Data extraction parameters were set as dynamic and non-linear iRT calibration with precision iRT was selected. Identification was performed using 1% precursor and protein q-value. iRT profiling was selected. Quantification was based on the peak areas of extracted ion chromatograms (XICs) of the 3-6 best fragment ions per precursor ion and q-value sparse data filtering was applied. Interference correction was selected and no normalization was applied. Differential protein abundance analysis was performed using a paired t-test and p-values were corrected for multiple testing, using the Storey method [82,83]. All identified and quantified protein groups were kept. Protein groups with q-value < 0.05 and absolute Log2(fold-change) > 0.5 were considered significantly altered (Supplementary Table 1).

For Experiment 2 and Experiment 4, DIA data were processed in Spectronaut v15 (15.1.210713.50606) using directDIA. Data was searched against the human proteome with 20,386 entries (UniProtKB-SwissProt), accessed on 07/31/2021. Trypsin/P was set as the digestion enzyme and two missed cleavages were allowed. Cysteine carbamidomethylation was set as a fixed modification while methionine oxidation and protein N-terminus acetylation were set as dynamic modifications. The Pulsar identification was performed using 1% PSM, peptide and protein group false discovery rate. The settings for the DIA analysis were the same as above. A quantity correction factor was applied to normalize the abundances to the cell counts. Differential protein abundance analysis was performed using a paired t-test, and p-values were corrected for multiple testing, using the Storey method [82,83]. Protein groups are required with at least two unique peptides. Protein groups with q-value < 0.05 and absolute Log2(fold-change) > 0.5 were considered significantly altered (Supplementary Tables 2 & 4).

For Experiment 3, DIA data were processed in Spectronaut v17 (17.6.230428.55965) using directDIA. Data was searched against the human proteome with 20,386 entries (UniProtKB-SwissProt), accessed on 07/31/2021. Trypsin/P was set as the digestion enzyme and two missed cleavages were allowed. Cysteine carbamidomethylation was set as a fixed modification while methionine oxidation and protein N-terminus acetylation were set as dynamic modifications. The Pulsar identification was performed using 1% PSM, peptide and protein group false discovery rate. The settings for the DIA analysis were the same as above. Differential protein abundance analysis was performed using an unpaired t-test, and p-values were corrected for multiple testing, using the Storey method [82,83]. Protein groups are required with at least two unique peptides. Protein groups with q-value < 0.05 and absolute Log2(fold-change) > 0.5 were considered significantly altered (Supplementary Table 3).

MetaVolcanoR package v.3.19 (https://github.com/csbl-usp/MetaVolcanoR/blob/master/) was used to integrate protein quantifications from 4 different MS experiments, which summarizes the differential expression p-values using Fisher’s method. Metafc= ‘mean’, and metahr =0.01 was used. Stiffness differential secretion of proteins of senescent cells vs. either proliferating or quiescent controls were done by ranking the fold change of senescent cells vs. control on either 2 kPa or plastic. Heatmap of protein fold changes of senescent vs. proliferating on either 2 kPa or plastic were graphed with Pheatmap v1.0.12 (https://cran.r-project.org/web/packages/pheatmap/index.html), and ordered by decreasing value.

### SA-β-Gal staining

SA-β-gal activity was determined using the Senescence Detection Kit (Abcam, Fremont, CA; #ab65351). Six days after doxorubicin or DMSO treatment (untreated), cells on each condition were fixed and stained for SA-β-gal as per the manufacturer’s protocol. Nuclei were stained using Hoechst 33342 (BD Pharmingen, San Diego, CA; #561908) and imaged using the Olympus IX70 microscope (Tokyo, Japan), followed by manual counting using ImageJ [84]. For each replicate, approximately 400-600 cells were counted.

### EdU assay

EdU staining was conducted using Click-iT EdU Alexa Fluor 488 HCS Assay (Thermo Fisher Scientific, Waltham, MA; #C10351). After doxorubicin treatment and 24 hours before collection, cells were changed into media containing 2.5 µM EdU. After 24 hours, cells were fixed and permeabilized and Click-iT Reaction Cocktail was added as per manufacturer’s protocol. Cells were incubated for 30 min in dark at RT and stained for Hoechst 33342 (BD Pharmingen, CA; #561908). Lastly, cells were imaged using Olympus IX70, approximately 500-800 cells were counted per replicate using Image J as described earlier.

### Bleomycin (BLM) induced IPF mouse model

C57BL/6N mice were purchased from Jackson Laboratory. All animal care protocols and procedures were performed in accordance with relevant guidelines and with approval by the Institutional Animal Care and Use Committee of the University Health Network (Toronto, Ontario, Canada). Six to eight weeks old mice were used for BLM injury. BLM 2.5 U/kg body weight was administered intratracheally as previously described [85]. Control mice received saline. Mice lungs were collected 14 days post-BLM administration.

### Immunohistochemistry

Mice lungs were fixed 48 hours in 10% buffered formalin prior to processing and Masson’s trichrome staining or immunohistochemistry with p21 primary antibody (Abcam, UK; # ab188224) (1:400 overnight at room temperature) and detection with ImmPRESS HRP Horse Anti-Rabbit IgG Polymer Detection Kit (Vector Laboratories, Newark, CA; # 7401). The tissue was imaged by light microscopy using a Leica DFC320 camera with the Leica application suite (LAS) software.

### RNA extraction and quantitative real-time PCR

Collected cell pellets were stored in TRIzol Reagent (Invitrogen, Waltham, MA; #15596026) for long-term storage at -80°C. Total RNA was prepared using the Zymo Research Direct-zol RNA Miniprep kit (Irvine, CA; #R2050). RNA was extracted as per the provided protocol and stored at -80°C for next-generation sequencing.

For other samples, RNA was reverse transcribed into cDNA using a High-Capacity cDNA Reverse Transcription Kit (Applied Biosystems, Foster City, CA; #4368813). Quantitative real-time PCR (qRT-PCR) reactions were performed as using the TaqMan Fast Advanced Master Mix (Applied Biosystems, Foster City, CA; #4444556) and CDKN1A TaqMan Gene Expression Assays Hs00355782_m1 (Applied Biosystems). Actin (ACTB) TaqMan Gene Expression Assay Hs99999903_m1 (Applied Biosystems) was used to control for cDNA quantity. qRT-PCR assays were performed on the QuantStudio 6 Pro Real-Time PCR System (Applied Biosystems).

### Statistical analysis and Data visualization

SA-β-gal, EdU and QPCR were measured using unpaired student T test.

Bar plots were visualized using GraphPad Prism 10.2.2 (La Jolla, CA). Box and dot plots were made with ggplot2 v3.5.1[86] and ggpubr v 0.6.0 (https://github.com/kassambara/ggpubr). Venn diagrams were plotted using ggvenn v.0.1.10 (https://github.com/yanlinlin82/ggvenn). Pathway and predicted downstream and upstream network were generated from QIAGEN Ingenuity Pathway Analysis (Venlo, Netherlands).

### Bulk RNA sequencing

RNA quantification, preparation of RNA library, and mRNA sequencing were conducted by Novogene Co., LTD (CA, US). About 20 million paired-end 150 bp reads per sample were generated from Illumina NovaSeq 6000 Sequencing System. FASTQ raw reads were analyzed using seqWorkflows (https://github.com/natmurad/seqWorkflows). Briefly, raw files were trimmed using trimmomatic, aligned using STAR and annotated using RSEM. Differential gene expression analysis between groups was done by DESeq2 R package (version 1.36.0) [87].

Genes with adjusted p-value < 0.05 and |log2(FoldChange)| > 0.5 were considered as significantly differentially expressed (Supplementary Table 9).

### Re-analysis of published single-cell RNA-seq data

These data were obtained from Gene Expression Omnibus (GEO) accessions GSE122960 [43], GSE128033 [44], GSE128169 [45], and GSE135893 [46]. The sample data from each study except GSE135893 were processed in Seurat (version 4.1.1) [88,89] as individual Seurat objects and filtered to include cells with <10% mitochondrial RNA and > 250 genes/features as a QC procedure. Raw RNA counts were normalized and stabilized using the SCTransform v2 function (SCT) and combined into one Seurat object. The R object from GSE135893 [46] which contained annotated cell clusters was directly imported into R and processed using the SCTransform function and later used as an anchor reference for the other two datasets. Seurat Transfer Anchors function was applied to the merged Seurat object to predict cell type in all samples. Cell type proportion (Supplementary Table 10) was calculated per cell type per study. Each of the four fibroblasts cell types, Myofibroblasts, Fibroblasts, PLIN2+ Fibroblasts, and HAS1 High Fibroblasts, were individually subsetted for downstream analysis. Differential gene expression between conditions were performed using FindMarker function with the parameters, min.pct=0.005 and logFC=0.1.

### Pathway analysis

Following differential expression analysis, Ingenuity Pathway Analysis (Qiagen, Netherlands) was used to identify enrichment in pathways for each comparison. DEGs and differential secreted proteins with Q-values < 0.05 and |Log2FC|> 0 were incorporated into the IPA canonical pathway analysis. IPA pathways that had a cut off of p < 0.05, z-score > |1.5| were plotted as results.

### Gene set enrichment score calculation and correlation analysis

Gene sets were selected from literature [9,53–55] and from the Molecular Signatures Database [49]. Details of selected gene sets are listed in Supplementary Table 6. Gene set enrichment scores were calculated using the Seurat AddModuleScore function. Significances were calculated with Wilcoxon Rank Sum test. Collagen-related gene set scores were measured by Gene Set Enrichment Analysis [49,90]. Pearson correlation was used to calculate correlation between gene sets using both healthy and ILD fibroblast cells.

## Data Availability

Raw data and complete MS data sets have been uploaded to the Center for Computational Mass Spectrometry, to the MassIVE repository at UCSD and can be downloaded using the following FTP link: ftp://MSV000095880@massive.ucsd.edu or via the MassIVE website: https://massive.ucsd.edu/ProteoSAFe/dataset.jsp?task=249d2196622f476baaa1b1394cd90c1f (MassIVE ID number: MSV000095880; ProteomeXchange ID: PXD055921).

The bulk RNA-sequencing data generated in this study are deposited in the GEO database under accession code [GSE282384].

The code for analysis will be available at GitHub [https://github.com/ZoeDUD/Stiffness_Modulate_Senescence].

## Acknowledgements

We thank Kristina Asoyan and Carmelita Honculada from University Health Network, Pathology Research Program (PRP) lab for technical assistance with Immunohistochemistry. We gratefully acknowledge the valuable support of Natália Faraj Murad for discussions on RNA-seq analysis.

## Funding

This work was funded, in part, through funds derived from the Buck Institute for Research on Aging (D.A.W.), the National Institutes of Health grant R01DK128435 (D.A.W.), the Huiying Memorial Foundation (D.A.W.), and Canadian Institutes of Health Research grant PJT-186165 (D.A.W.). The project was also funded in part through the University of Southern California and Buck Institute Nathan Shock Center (P30 AG068345). This work was supported by the NIH-OD shared instrumentation grants S10 OD028654 (B.S.) for the Orbitrap Eclipse Tribrid mass spectrometry system and S10 OD016281 (Buck Institute) for the TripleTOF 6600 mass spectrometry system. T.R.V. was funded by a T32 NIH fellowship grant (NIA T32 AG000266).

## Author information

H.D. contributed to the experimental design, execution, and analysis of the study. H.D. and D.A.W. drafted the manuscript. J.P.R., J.B., N.B., J.B.B. and B.S. performed mass spectrometry, data processing and analysis. J.P.R, T.R.V., M.M., and P.M. assisted with *in vitro* experiments and edited the manuscript. L.G. and A.N. provided BLM mice samples. F.W. and D.F. assisted bioinformatic analysis. S.W., L.G., N.C. and V.C. assisted with IHC. D.W., B.S. and J.C. supervised the study. All authors had the opportunity to view and edit the manuscript.

## Competing interests

D.F. and D.A.W. are co-founders of Cosmica Biosciences, a company that studies altered biological aging under changes in mechanical force.

## Notes

### Summary of Updates

The method is updated with GEO depository and GitHub information.

