## Supplementary Figures for "Substrate Stiffness Dictates Unique Doxorubicin-induced Senescence-associated Secretory Phenotypes and Transcriptomic Signatures in Human Pulmonary Fibroblasts"

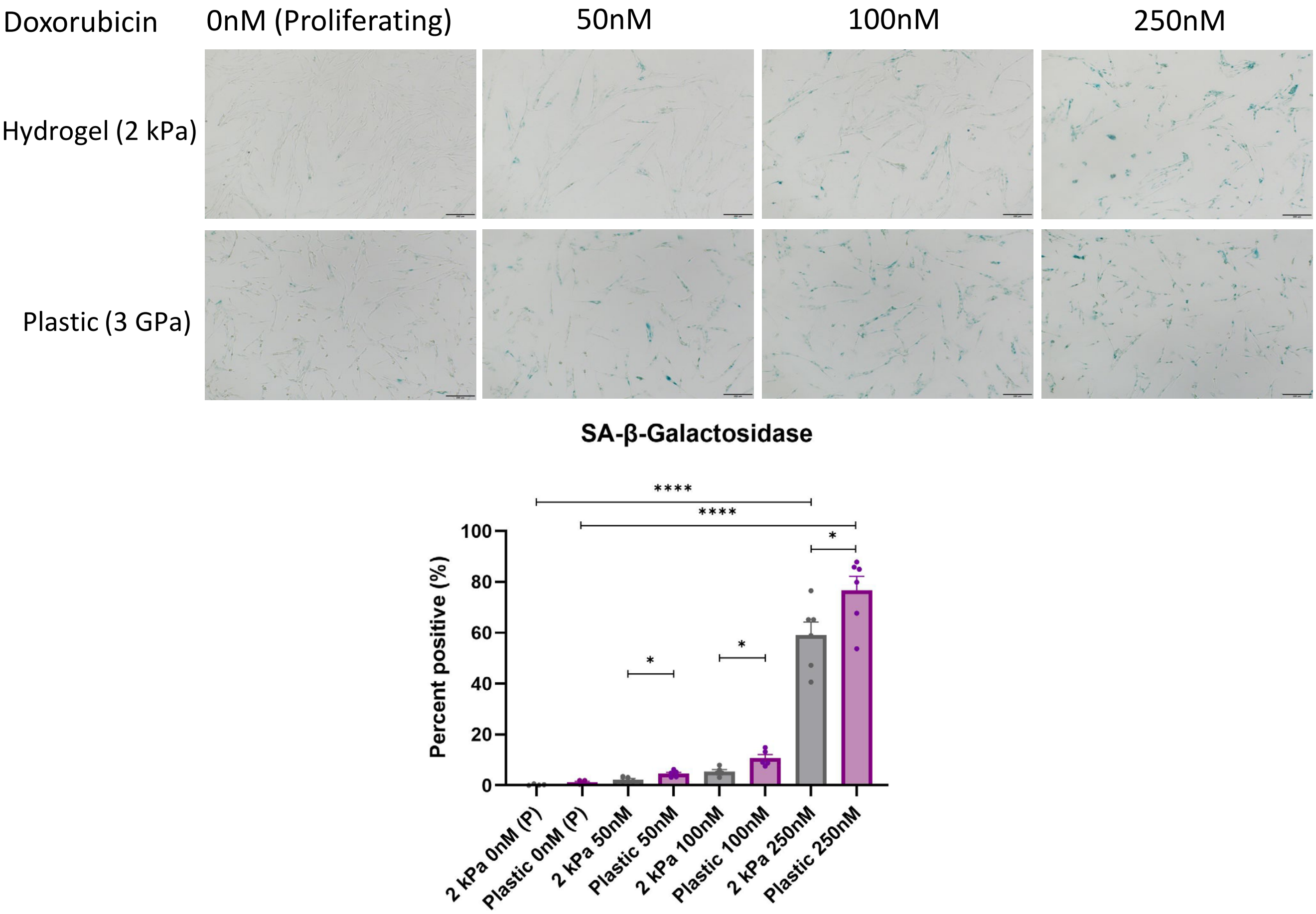


Supplementary Figure 1. Representative images of SA-β-Gal staining and quantification across different doxorubicin(doxo) doses. Error bars represent SEM of biological variation, statistics evaluated via unpaired student T test, data presented as biological replicate indicated by dots (n = 4 for untreated controls, n=5 for 50nM and 100nM doxo treated conditions, and n=6 for 250nM doxo treated conditions; ****P <0.0001, *P < 0.05).

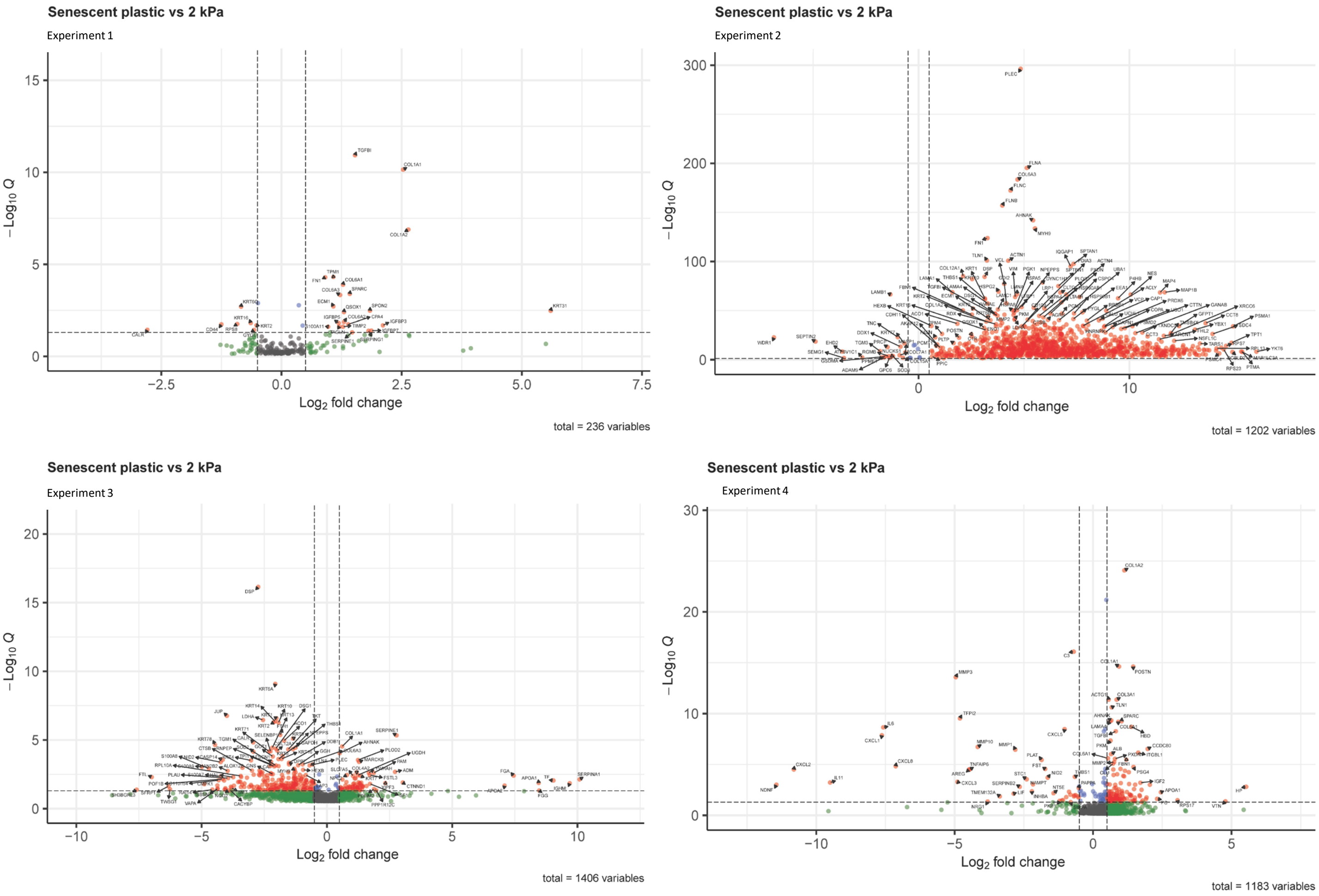


Supplementary Figure 2. Volcano plot of four independent mass spectrometry experiments (1-4) showing proteins secreted into the culture supernatant by senescent IMR-90 fibroblasts treated with 250nM doxorubicin cultured on plastic plate vs. 2 kPa hydrogel (n= 4 for MS1, n= 6 for MS2 and MS3, n=3-4 for MS4; volcano cut off is Q < 0.05, Log2FC > |0.5|.

A

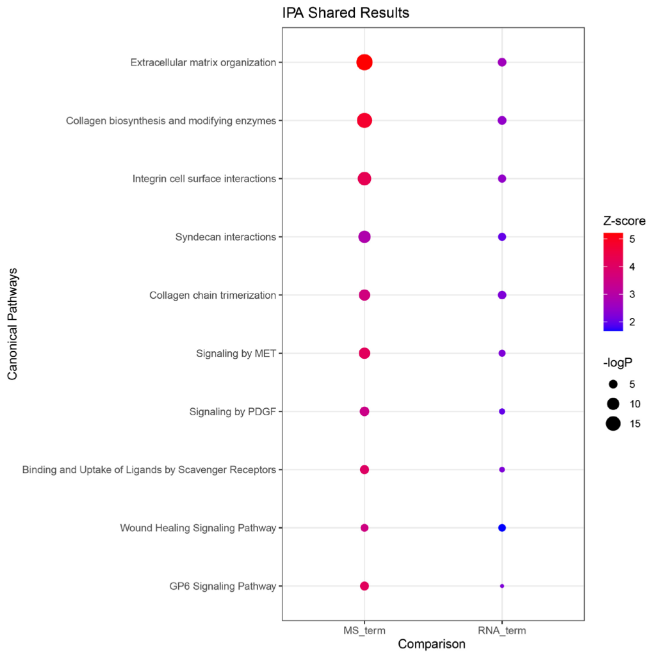


B

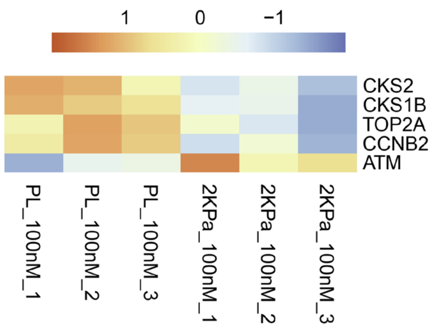


Supplementary Figure 3. Overlapping proteomic and transcriptomic signatures show that stiffness drives changes in senescent IMR-90 fibroblasts.

A. Pathway enrichment (IPA) of senescent IMR-90 fibroblasts treated with 250nM doxo cultured on plastic vs. 2 kPa. The proteomic MS result and transcriptomic RNA result were placed side by side for comparison.
B. Heatmap of gene expression that contributes to G2/M damage checkpoint regulation pathway.

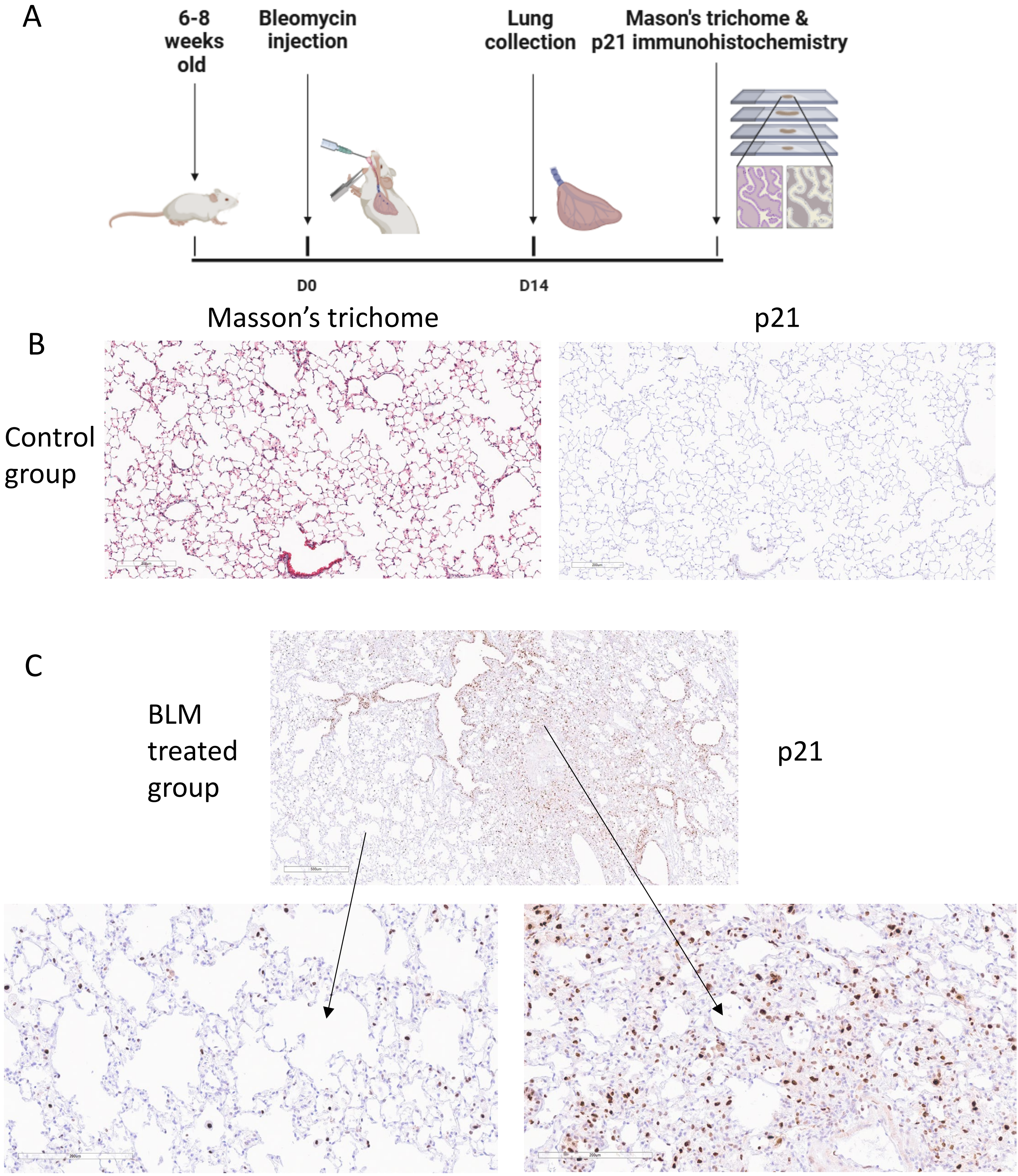


Supplementary Figure 4. Bleomycin-induced (BLM) fibrosis mice lungs showed clusters of p21 positive cells in areas of fibrosis.

a. Schematic diagram of IPF mouse model.
b. Saline treated control mouse lung with Masson's trichome (left) and p21 (right) staining.
c. Different regions of BLM treated mouse lung show different level of fibrosis and alveolar architectural distortion and varying levels of p21 staining, with the right region showing fibrosis and clusters of p21.

A

| | GSE128033 | GSE128169 | GSE122960 | GSE135893 |
|---|---|---|---|---|
| Sample # | 18 | 11 | 17 | 20 |
| Healthy Control # | 10 | 1 | 8 | 10 |
| ILD # | 8 | 10 | 9 | 20 |

B

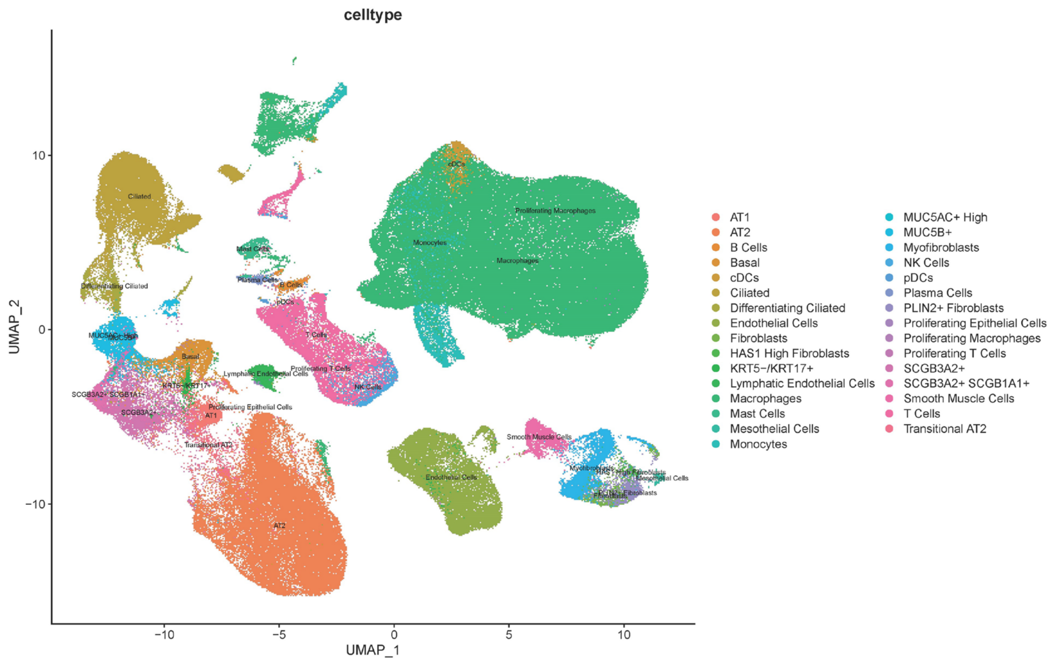


C

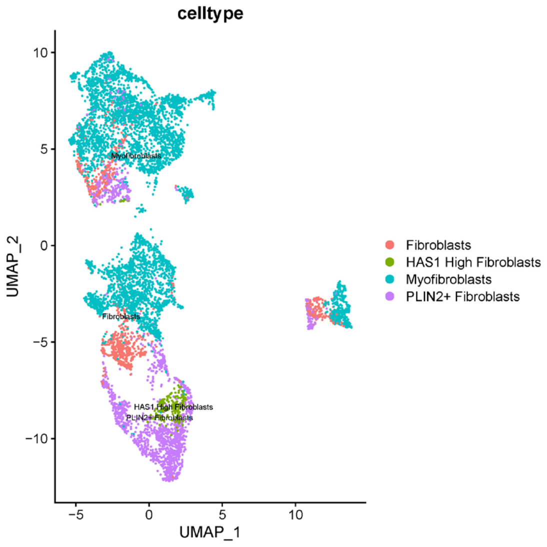


Supplementary Figure 5. Reanalysis of published human lung single-cell RNA sequencing data.

a. Metadata of 4 published single-cell RNA sequencing data that was used in this study.
b. Uniform Manifold Approximation and Projection (UMAP) of 66 human lung samples identified 31 cell types.
c. UMAP of only fibroblast cells from 66 samples.

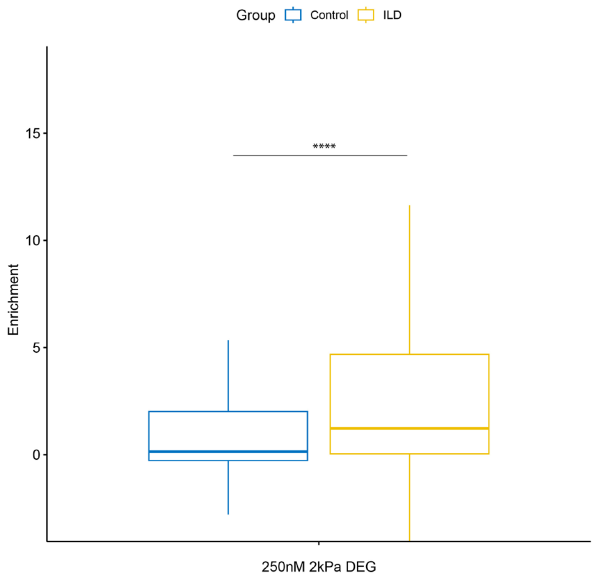


Supplementary Figure 6. Gene set enrichment score of ILD and control fibroblasts using the DEGs from 250nM doxo-induced senescent cells cultured on 2kPa hydrogel compared to plastic.

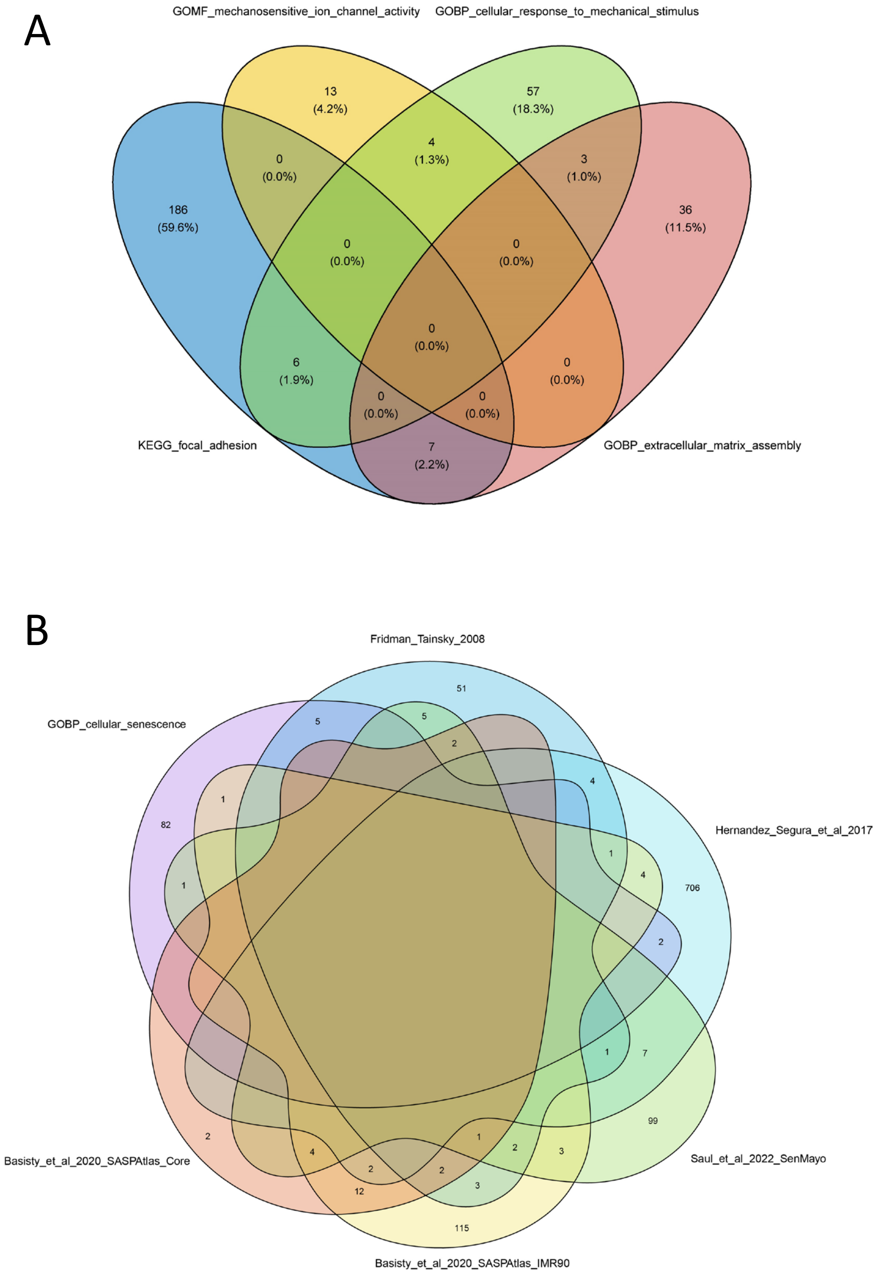


Supplementary Figure 7. Venn diagram of overlapping genes and proteins between mechanotransduction-related and senescence signatures selected from literature

a. Genes that overlap between selected mechanotransduction-related gene sets.

b. The overlap between genes and proteins in selected cellular senescence signatures.
